# Diffusion-based stimulus optimization reveals functional organization across higher visual cortex

**DOI:** 10.64898/2026.05.12.724119

**Authors:** Margaret M. Henderson, Andrew F. Luo, Sungjoon Park, Michael J. Tarr, Leila Wehbe

## Abstract

Characterizing the fine-grained functional organization of human higher visual cortex remains a central challenge, as traditional neuroimaging experiments constrain the diversity of stimuli that can be sampled. In prior work we addressed this challenge by developing a novel data-driven tool, termed “BrainDiVE” (Luo et al. 2023), which synthesizes naturalistic images predicted to strongly activate specific brain regions. BrainDiVE leverages pretrained image diffusion models guided by gradients from an image-computable fMRI encoding model. Here, we experimentally validated BrainDiVE by generating images predicted to maximally activate different functional regions of interest and then presenting them to new participants (n=12) in an fMRI study. The model-generated images elicited robust, spatially specific responses in the targeted brain regions, producing significantly greater category selectivity than natural images, validating the method’s ability to capture generalizable neural tuning properties in human ventral visual cortex. We further showed that region-targeted images exaggerate specific sets of low-level and mid-level image statistics, suggesting that category-selective regions are tuned to continuous directions in feature space. Moreover, we demonstrated fine-grained experimental control by differentially activating two face-selective regions, the occipital face area (OFA) and fusiform face area (FFA), providing additional evidence that these regions encode distinct aspects of faces. Finally, we identify a posterior-to-anterior functional gradient within the occipital place area (OPA), suggesting topographic organization based on scene properties such as distance and indoor-outdoor location. These findings enhance our understanding of the representational structure of category-selective regions and introduce a new paradigm for probing neural selectivity in human visual cortex.

## Introduction

A central goal of visual neuroscience research is to characterize the response properties of cortical populations (Tanaka, 1996) with the aim of understanding their functional role in perception. Recent advances in machine learning, including deep neural networks, evolutionary optimization, and generative image models, have enabled continuous synthesis of stimuli optimized for specific neural populations or brain regions (Bashivan et al., 2019; Cowley et al., 2026; Gu et al., 2022; Ponce et al., 2019; Ratan Murty et al., 2021; Walker et al., 2019). These approaches generate images that match or exceed the responses evoked by natural images, yielding supra-activating stimuli that can provide data-driven insight into the fine-grained selectivity of cortical populations.

Similar applications of generative models to population data from higher visual cortex–particularly for human fMRI–have been relatively rare. The typical paradigm for inferring neural tuning properties from region-optimized stimuli is a closed-loop in which the empirical response to a region-optimized stimulus is measured, a necessary step to validate that the generated stimulus evokes the expected level of activity. Such closed loop paradigms have been successfully used to control population activity in primary visual cortex (Walker et al., 2019), V4 (Bashivan et al., 2019; Cowley et al., 2026) and portions of IT cortex (Ponce et al., 2019) via electrophysiological recordings. In other recent work, synthesized images were used to “parametrically control” IT responses along model-derived encoding axes, enabling a finer-grained understanding of model-brain alignment (Prince et al., 2025). However, it is not clear if closed-loop methods can be applied to population-level functional magnetic resonance imaging (fMRI) signals measured in the human brain. Previous work in which region-optimized stimuli have been selected or generated based on fMRI signals has produced mixed results. It remains unclear whether such synthetic stimuli reliably elicit greater neural activation than natural images (Gu et al., 2022, 2023; Leeds & Tarr, 2016). This leaves an open question that recent advances in generative modeling are well positioned to address. A positive outcome would have important implications: if fMRI activation patterns in human visual cortex can be systematically controlled using region-optimized images, this would open up new avenues for investigating the organization of higher visual cortex and clarify how visual and semantic properties contribute to its structure.

To this end, we developed BrainDiVE (Brain Diffusion for Visual Exploration; Luo et al., 2023), a generative framework that produces synthetic images predicted to maximally activate a cortical region of interest. BrainDiVE leverages a large pre-trained latent image diffusion model (LDM; Rombach et al., 2022), along with a highly accurate forward encoding model (Naselaris et al., 2011; Serences & Saproo, 2012; Wang et al., 2023) that predicts fMRI activation given an input image. By perturbing the diffusion synthesis procedure in the direction of the encoding model gradients, BrainDiVE can synthesize “optimal” images for a given population of voxels or region of higher visual cortex. Validating this approach, the resulting BrainDiVE synthesized images contain content that aligns with the expected semantic selectivity of category-selective cortical regions of interest (ROIs;Figure 1). Differences in synthesized BrainDiVE images further dissociate the response properties of functionally related ROIs, such as the occipital face area (OFA) and the fusiform face area (FFA), as well as subregions within a single ROI, such as the occipital place area (OPA).

**Figure 1.**
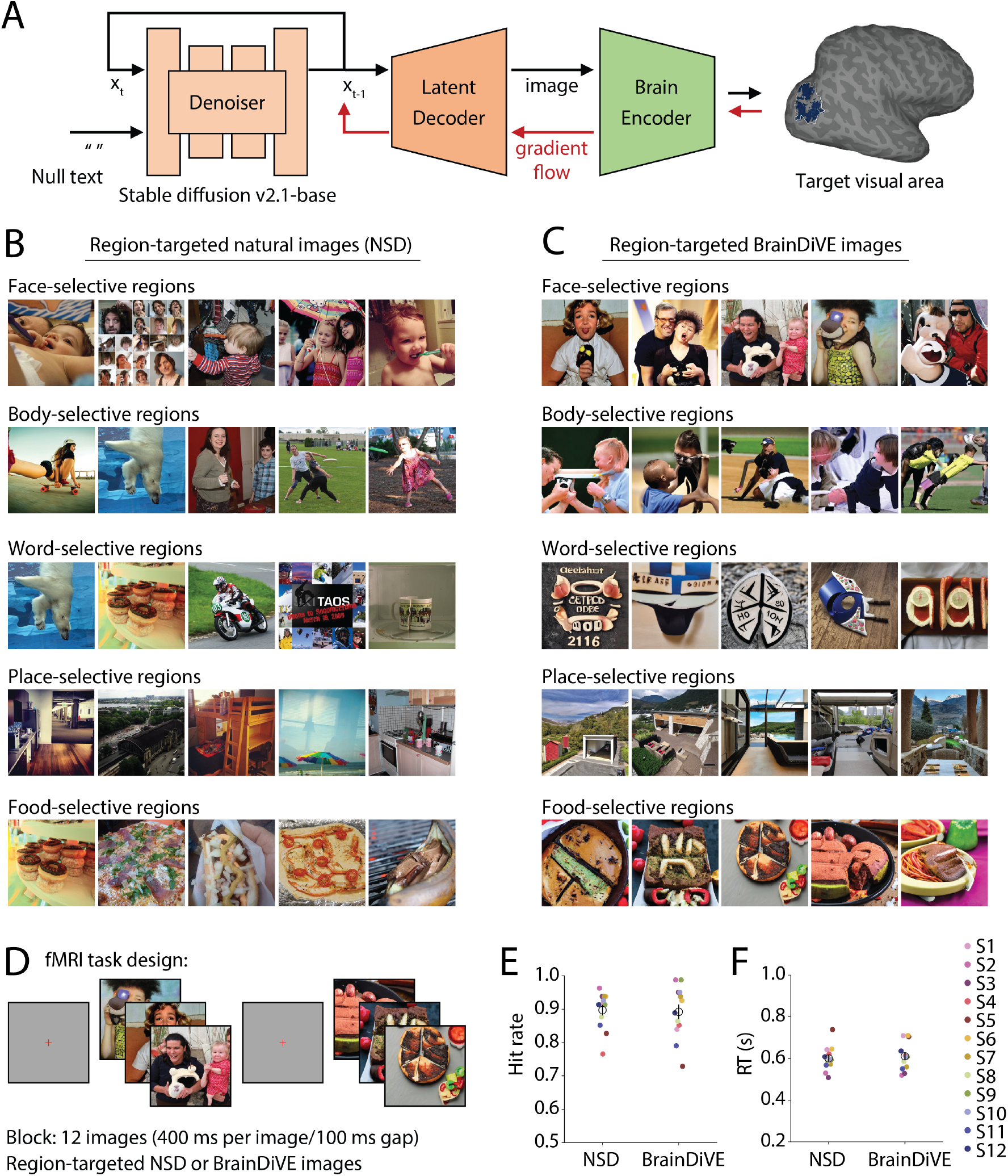
Method for generating region-optimized stimuli and overview of fMRI experiment. **(A)** Schematic of the BrainDiVE image optimization procedure (Luo et al., 2023). We leverage a pretrained latent diffusion model, which is trained to synthesize natural images by iterative denoising (Rombach et al., 2022). We also construct an encoding model, which is trained to map images to cortical responses, leveraging features from a deep neural network (CLIP ViT-B/16; see *Methods* for details), fitted to neural data from the Natural Scenes Dataset (NSD; Allen et al., 2022). During denoising, we perturb the LDM latent vector in a direction predicted to maximize activation in a target brain region. We define a brain-maximization objective and use backpropagation to compute its gradient with respect to the latent vector (see *Methods* for details). **(B-C)** Example images targeting category-selective regions for NSD participant S1; **(B)** The five top-activating images for each ROI in NSD (i.e., natural scenes viewed during fMRI); **(C)** The top-5 images generated using BrainDiVE (see *Methods*); **(D)** Schematic of behavioral tasks performed in the scanner. On alternating scan runs, participants viewed either BrainDiVE-optimized images targeting specific ROIs (BrainDiVE runs) or natural scene images that elicited maximal activation in those same ROIs in NSD participants (NSD runs; see *Methods*). Stimuli were shown in a block design (12 images per block, 400 ms per image with 100 ms gap between images), while participants performed a one-back repeated image detection task; **(E-F)** Behavioral performance for BrainDiVE and NSD runs; **(E)** Hit rate–the proportion of correctly detected image repeats–on the one-back task performed in the scanner; **(F)** Response time (RT). Colored dots indicate individual participants, open circles and error bars indicate mean and SEM across participants (n=12). All photographs, including those depicting people, are from the Microsoft COCO dataset (Lin et al., 2014) and are used in accordance with the COCO dataset’s Creative Commons Attribution 4.0 license.

Here, we test the prediction that BrainDiVE-generated, region-optimized images elicit stronger neural responses in target ROIs than the highest-activating natural scene images for those same areas. Because BrainDiVE operates offline, this prediction has not yet been directly tested. To address this, we conducted an fMRI experiment in which 12 participants viewed BrainDiVE-generated images and natural images selected to target specific higher visual regions as established in an independent cohort (original participants from the Natural Scenes Dataset;NSD; Allen et al., 2022). We examined voxel-wise responses across visual cortex, as well as functionally-localized, category-selective ROIs. Previewing our results, we find that category-selective responses are reliably larger in magnitude for BrainDiVE region-optimized stimuli than for region-targeted natural stimuli. We further provide quantitative analyses of the generated images, yielding new insights into the stimulus properties that most strongly drive category-selective responses in human higher visual cortex.

## Results

### Generating region-targeted experimental stimuli

To characterize the pattern of functional selectivity across human higher visual cortex, we used Brain-DiVE (Luo et al., 2023) to synthesize stimulus images optimized to yield maximal activation in target visual cortex ROIs based on whole-brain 7T fMRI data from participants viewing complex natural scene images (data from NSD; Allen et al., 2022). BrainDiVE combines a large pretrained latent diffusion model (LDM; Rombach et al., 2022) with a voxel-wise encoding model based on features extracted from a deep neural network (Naselaris et al., 2011; Radford et al., 2021) to generate complex, naturalistic images that capture the preferred stimulus properties of target cortical regions (Figure 1; see *Methods* for more details).

When images are optimized to target category-selective regions (i.e., face-, body-, place-, word-, food-selective regions) in individual NSD participants, BrainDiVE reliably generates images that reflect the expected category-specific content for each targeted region. More specifically, as shown in Figure 1C, images targeted to face-selective regions contain recognizable human faces, images targeted to body-selective regions contain recognizable human forms and actions, images targeted to place-selective regions contain recognizable indoor and outdoor scenes, images targeted to word-selective regions contain features that resemble written characters, and images targeted to food-selective regions contain objects and materials that are recognizable as food.

As shown in Figure 1B, similar category-level differences can be observed by selecting, from the natural images viewed by NSD participants, those that elicited the highest activation in the same targeted visual regions. Importantly, because BrainDiVE images are not constrained to a fixed set of experimental stimuli, they can more precisely and continuously capture the image properties that drive each target region. Consistent with this point, BrainDiVE images are more semantically aligned with the expected target category of each ROI as compared to the NSD images. For example, whereas the top-activating NSD images for word-selective voxels span a mixture of categories, including food and motor vehicles, BrainDiVE images for the same regions contain a larger proportion of written characters. We previously quantified these differences using a CLIP-based semantic probing method (Luo et al., 2023), demonstrating greater semantic specificity for BrainDiVE-generated images relative to top-activating natural images. These results suggest that BrainDiVE effectively disentangles the properties of complex natural scenes that drive individual category-selective ROIs in visual cortex. Moreover, BrainDiVE captures functional differences in selectivity both between individual ROIs within the face-processing network (OFA, FFA;Figure 5A) and within individual ROIs, such as the occipital place area (OPA;Figure 6), revealing functionally distinct voxel clusters and demonstrating sensitivity to fine-grained functional organization.

In the present study we tested whether BrainDiVE images (1) represent a distillation of the most important visual features for driving a given neural population; (2) generalize to a new set of participants whose data were not used in the BrainDiVE synthesis process. We ran an fMRI experiment in which 12 participants were scanned while viewing BrainDiVE-generated images targeted to different category-selective regions. As a baseline, participants also viewed the NSD images that elicited the highest activation in the same category-selective regions in the original NSD cohort. Experimental images (BrainDiVE and NSD) included 9 total conditions. In the first 5, images were targeted to broad networks of category-selective voxels, defined by combining ROIs with shared category selectivity (face-, body-, place-, word-, and food-selective ROIs; see *Methods*). To enable more fine-grained functional comparisons, the remaining 4 conditions included images targeted for OFA and FFA individually, and for two subregions of OPA that we functionally identified in NSD participants (Luo et al., 2023; see *Methods*). In separate scan runs, participants viewed BrainDiVE or NSD images in a block design while performing a one-back repetition detection task. No behavioral task performance differences were observed between NSD and BrainDiVE runs, either in accuracy (hit rate: NSD: 0.90 ±0.05; mean ±SD;BrainDiVE: 0.89 ±0.08; two-sided paired *t*-test with permutation: *t* (11) = 0.32, *p* = 0.756) or response time (RT: NSD: 600.58 ±56.92 s;BrainDiVE: 610.78 ±65.36 s; two-tailed paired *t*-test with permutation: *t* (11) = −1.12, *p* = 0.293).

### Selective responses of category-selective networks to region-targeted stimuli

Our primary hypothesis was that BrainDiVE-generated images targeted to particular sets of category-selective regions would yield greater fMRI activation than natural images targeted to the same regions, with this increase in activation being specific to the targeted regions. This effect was expected to be observed both for broad category-selective networks of regions, defined using independent functional localizer scans, and for more fine-grained regions (OFA vs. FFA, and functional clusters within OPA). We first discuss results with respect to the broad category-selective networks, and then turn our attention to more fine-grained patterns of selectivity.

To evaluate whether BrainDiVE images elicited greater category-selective responses than natural images in category-selective ROIs, we computed a *t*-statistic contrasting responses to images targeting one set of category-selective ROIs versus others. As predicted, BrainDiVE images resulted in larger *t*-statistics than NSD images in the targeted regions (Figure 2). As shown in Figure 2A, voxels in and around the face-selective regions show positive *t*-statistics for the contrast of face-targeted versus other images for both the NSD (top) and BrainDiVE (bottom) conditions. The magnitudes of these *t*-statistics are greater for BrainDiVE images than for NSD images. Similar effects are observed for body-selective (Figure 2B; EBA), word-selective (Figure 2C; VWFA), place-selective (Figure 2D; OPA, PPA) and food-selective regions (Figure 2E; Food ROI). Interestingly, contrasts also revealed negative *t*-statistics in voxels outside the targeted regions, reflecting selective, differential activation by each group of images for different cortical regions. These negative *t*-statistics were often greater in magnitude for BrainDiVE images versus NSD images. For example, comparing BrainDiVE versus NSD images, voxels in PPA and food-selective ROIs showed more negative *t*-statistics when comparing face-targeted versus other images (Figure 2A). These results indicate that BrainDiVE images, compared to natural images, elicit more spatially specific category-selective responses across ventral and lateral visual cortex.

**Figure 2.**
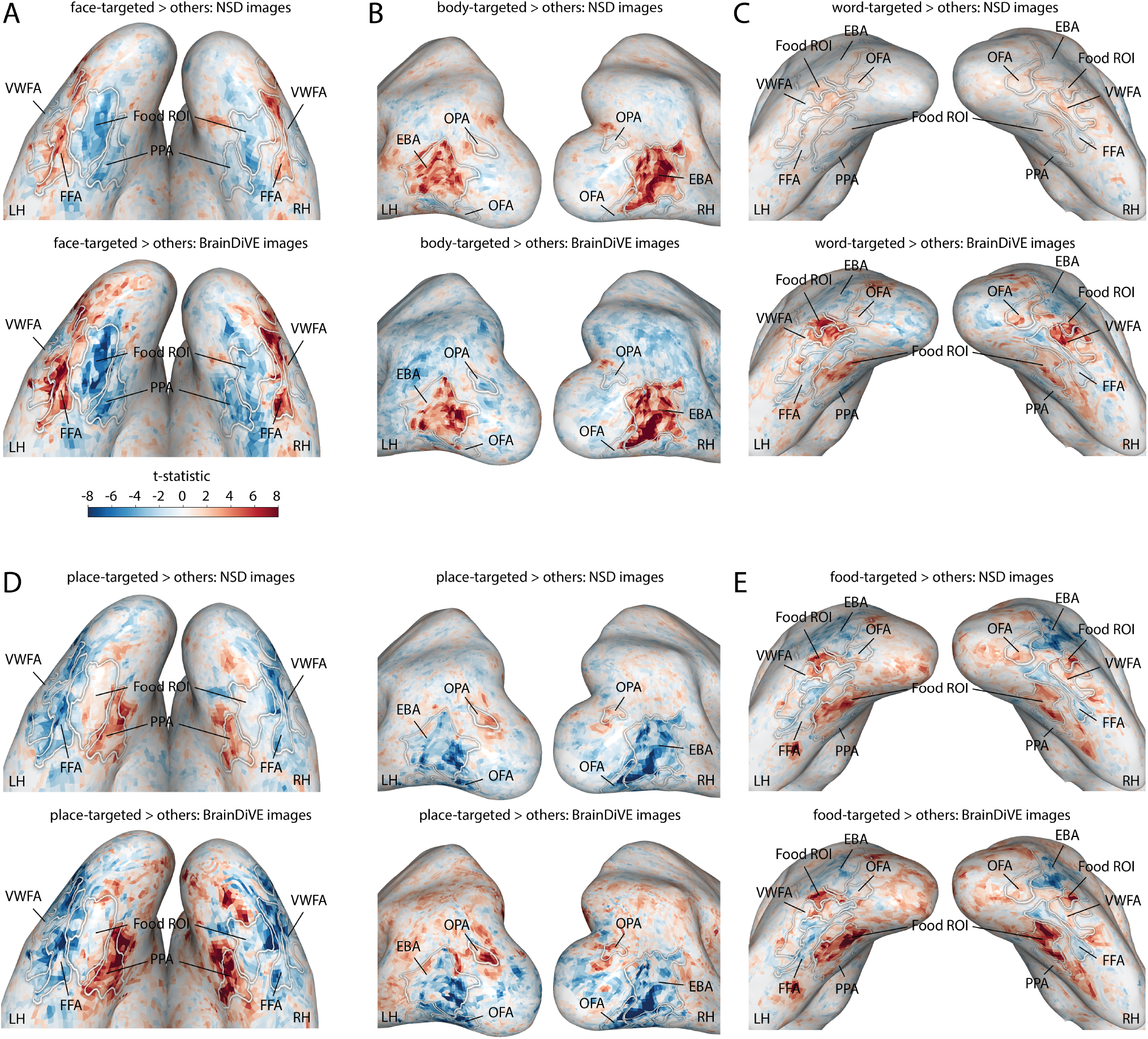
Category-selective responses elicited by BrainDiVE and NSD images. Data shown are for one participant (S03), projected onto an inflated cortical surface. Each plot shows *t*-statistics that reflect a contrast between images targeting a given set of category-selective ROIs as defined in the NSD participants (e.g., face-selective), versus images targeting all other ROIs. For all contrasts, the set of conditions compared is: face-targeted, body-targeted, word-targeted, place-targeted, food-targeted, where “other” is defined as the four conditions that were not the category of interest. Selectivity for **(A)** face-targeted; **(B)** body-targeted; **(C)** word-targeted; **(D)** place-targeted; and **(E)** food-targeted images. White labeled outlines indicate ROIs defined using an independent functional localizer task; see *Methods*.

Quantifying these effects at the level of functionally-defined ROIs reveals that they are consistent across participants (Figure 3). For each ROI (FFA, OFA, EBA, VWFA, PPA, OPA, and food-selective), we computed the *t*-statistic contrasting responses to images targeting the ROI’s corresponding category versus other categories, averaging these values across voxels within the ROI. All ROIs showed positive average *t*-statistics for both NSD images and BrainDiVE images, reflecting larger average responses to images that were targeted to that ROI versus images targeted to other ROIs (Figure 3A-E). Consistent with the hypothesis that BrainDiVE distills the features driving ROI selectivity, these positive *t*-statistics were higher for BrainDiVE images than for NSD images for many ROIs. Across all ROIs, a two-way repeated measures ANOVA revealed a significant main effect of condition (NSD vs. BrainDiVE), as well as a significant main effect of ROI and an interaction between ROI and condition (ROI: *F* (6, 66) = 14.50, *p <* 0.0001;Condition: *F* (1, 11) = 98.93, *p <* 0.0001;ROI:Condition: *F* (6, 66) = 5.25, *p <* 0.0001; all *p*-values obtained using permutation test; see *Methods*). Posthoc tests revealed that *t*-statistics for BrainDiVE images were significantly greater than for NSD images in FFA, OFA, EBA, VWFA, PPA, and OPA individually (BrainDiVE *t* vs. NSD *t*; two-tailed paired *t*-test with permutation;FFA: *t* (11) = 5.82, *p* = 0.001;OFA: *t* (11) = 3.55, *p* = 0.007;EBA: *t* (11) = 4.04, *p* = 0.003;VWFA: *t* (11) = 7.88, *p* = *<* 0.0001;PPA: 8.02, *p <* 0.0001;OPA: *t* (11) = 3.84, *p* = 0.006). This effect was highly consistent across participants, with FFA, VWFA, and PPA showing the expected pattern in all 12 participants (BrainDiVE *t >* NSD *t* for 12/12 participants in FFA, VWFA, PPA, 9/12 participants in OFA, 11/12 participants in EBA, 10/12 participants in OPA).

**Figure 3.**
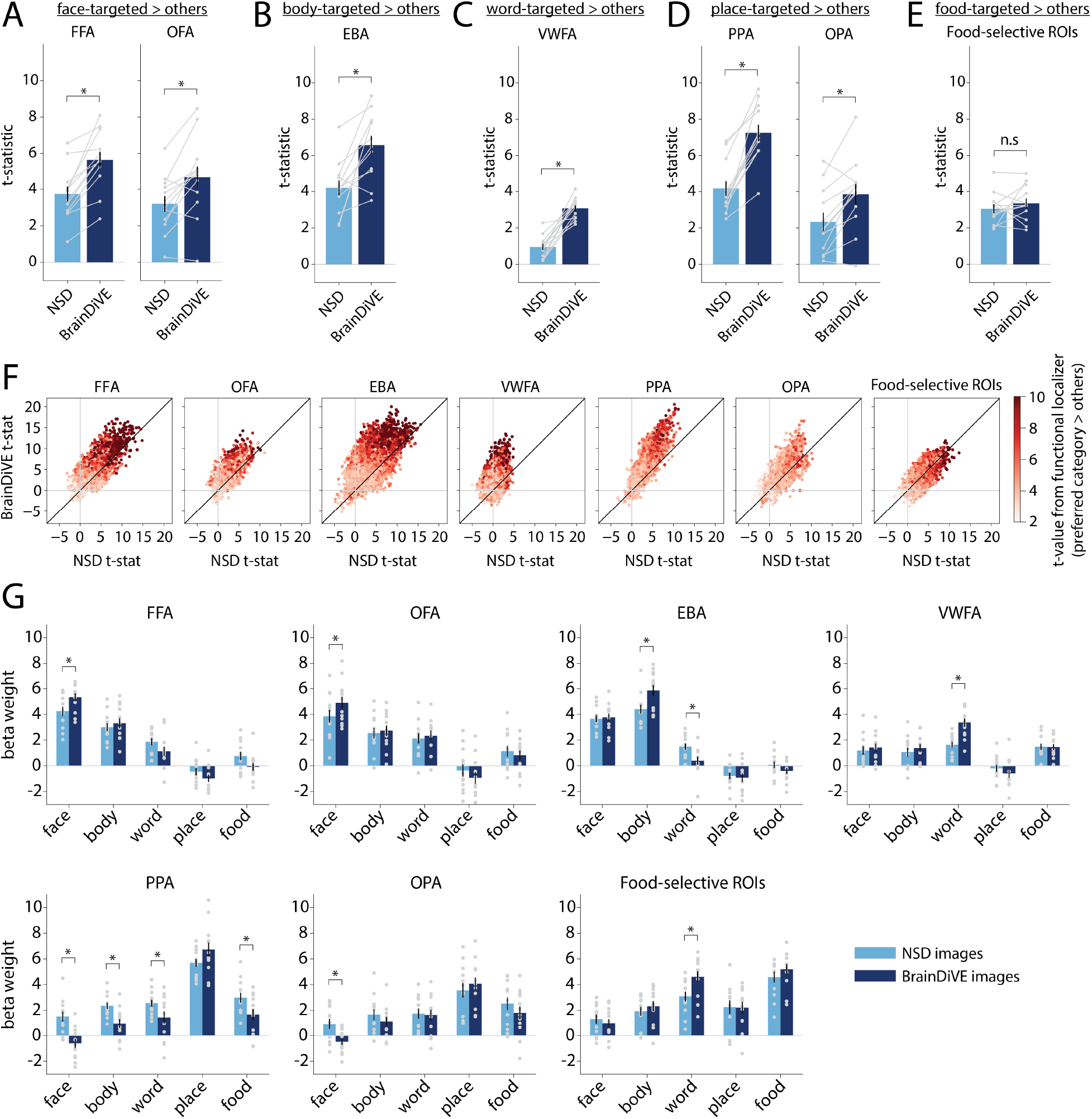
Comparison of category-selective responses elicited by BrainDiVE and NSD images, across all participants. **(A-E)** Average *t*-statistics for each ROI and each condition, where *t*-statistics reflect a contrast between images that were targeted to a specified set of category-selective ROIs versus images that were targeted to other ROIs. Contrasts capture selectivity for **(A)** face-targeted; **(B)** body-targeted; **(C)** word-targeted; **(D)** place-targeted; and **(E)** food-targeted images. Bar heights and error bars reflect the mean and SEM across participants (n=12); light gray dots and lines indicate single participant data, which is averaged across voxels within each ROI. Asterisks over pairs of bars indicate a significant difference (paired *t*-test with permutation, all *p <* 0.01, see text for test values). **(F)** Scatter plots showing the relationship between *t*-statistics computed for BrainDiVE versus NSD images. Each point represents a single voxel, voxels are pooled across all participants. Dot color indicates the *t*-statistic for that voxel from an independent functional localizer, capturing selectivity for the preferred category (e.g., for FFA, this value reflects face selectivity as measured in the functional localizer). The diagonal indicates a slope of 1. **(G)** Beta weights associated with each individual image category, for BrainDiVE and NSD images. Each panel represents one ROI; values shown reflect the beta weights associated with one image type (e.g., “face” refers to images targeted to face-selective ROIs in NSD participants). Different colors indicate beta weights from NSD or BrainDiVE runs. Bar heights and error bars indicate mean and SEM across participants (n=12); gray dots represent individual participants. Asterisks above pairs of bars indicate a significant difference between BrainDiVE and NSD (paired *t*-test with permutation; all *p <* 0.01; see Supplementary Table 2 for test statistic values).

Food-selective ROIs, however, showed a different pattern from the other ROIs. Although these ROIs showed reliable selectivity, responding more strongly to food-targeted images than to other images, this selectivity did not differ significantly between BrainDiVE and NSD images (BrainDiVE *t* vs. NSD *t*; two-tailed paired *t*-test with permutation; *t* (11) = 0.98, *p* = 0.363). Because the food-selective ROI consisted of non-adjacent medial and lateral patches (see *Methods* and Jain et al., 2023), we tested whether responses to BrainDiVE images differed between these two subregions (Supplementary Figure 1). Similar effects were observed across the medial and lateral patches, with neither subregion showing a significant difference in food selectivity between NSD and BrainDiVE images.

To understand how these effects varied across voxels within each ROI, we plotted the distribution of single-voxel *t*-statistics for BrainDiVE versus NSD runs as scatter plots (Figure 3F). For most ROIs, slopes were steep (slope *>* 1) with many points above the diagonal, indicating greater category-selectivity when measured using BrainDiVE. Moreover, voxels further above the diagonal tended to exhibit greater selectivity in an independent category localizer (higher *t*). This indicates that the largest BrainDiVE-related increases in selectivity occur in voxels that are also highly selective during the functional localizer task, suggesting that BrainDiVE images preferentially target voxels in proportion to their degree of category selectivity.

The increase in category selectivity for BrainDiVE images, as measured by *t*-statistics, could arise from several mechanisms. One possibility is that BrainDiVE images more effectively drive the regions they target, increasing “on-target” responses relative to NSD images. A second possibility is that BrainDiVE images reduce responses in non-target regions, decreasing “off-target” responses relative to NSD images. To distinguish between these possibilities, we examined each region’s response profile across category conditions (i.e., category-specific beta weights). As shown in Figure 3G, these profiles differed across ROIs. For face-selective, body-selective, and word-selective ROIs, BrainDiVE images led to increased on-target responses: FFA and OFA showed significantly higher responses to face-targeted BrainDiVE versus NSD images (two-tailed paired *t*-test with permutation; *p <* 0.01; see Supplementary Table 2 for all test values), EBA showed significantly higher responses to body-targeted BrainDiVE versus NSD images, and VWFA showed significantly higher responses to word-targeted BrainDiVE versus NSD images. In addition to these increased on-target responses, EBA showed significantly smaller responses to word-targeted BrainDiVE versus NSD images. In contrast, in place-selective regions PPA and OPA, the difference between BrainDiVE and NSD appeared to be driven by a reduction in off-target responses. PPA showed significantly smaller responses to face-targeted, body-targeted, word-targeted, and food-targeted BrainDiVE versus NSD images, but a non-significant difference between place-targeted BrainDiVE versus NSD images. Similarly, OPA did not show a significant difference between place-targeted BrainDiVE versus NSD images, but did show significantly smaller responses to face-targeted BrainDiVE versus NSD images. As in our other analyses, food-selective ROIs showed a different pattern from the other category-selective regions. Food-selective ROIs had a numerically greater but nonsignificant difference between food-targeted BrainDiVE versus NSD images, but did show significantly greater responses to word-targeted BrainDiVE versus NSD images. This result may be partially explained by overlap between food-selective and word-selective regions in the NSD dataset; we discuss this further in the *Discussion*.

### Image statistics of region-targeted stimuli

Given that BrainDiVE region-optimized images elicited significantly stronger responses in multiple category-selective ROIs, we next examined the image properties underlying this difference. In prior work (Luo et al., 2023), we showed that BrainDiVE images differ from NSD images in their semantic properties: using a zeroshot CLIP classifier (Radford et al., 2021), BrainDiVE images were more frequently classified as belonging to the expected category (e.g., face-targeted images classified as faces) as compared to region-targeted NSD images. This suggests that BrainDiVE images may capture a distilled representation of a region’s preferred category in semantic space.

Here, we evaluate whether low-level and mid-level image statistics contribute to this effect. For each stimulus image, we quantified the average luminance, contrast, color statistics, edge curvature, and spectral statistics (Figure 4 and Supplementary Figure 2; see *Methods* for details). For comparison, we computed the same set of statistics for 1,000 randomly-selected NSD images and 1,000 random LDM-generated images without brain guidance (see *Methods*).

**Figure 4.**
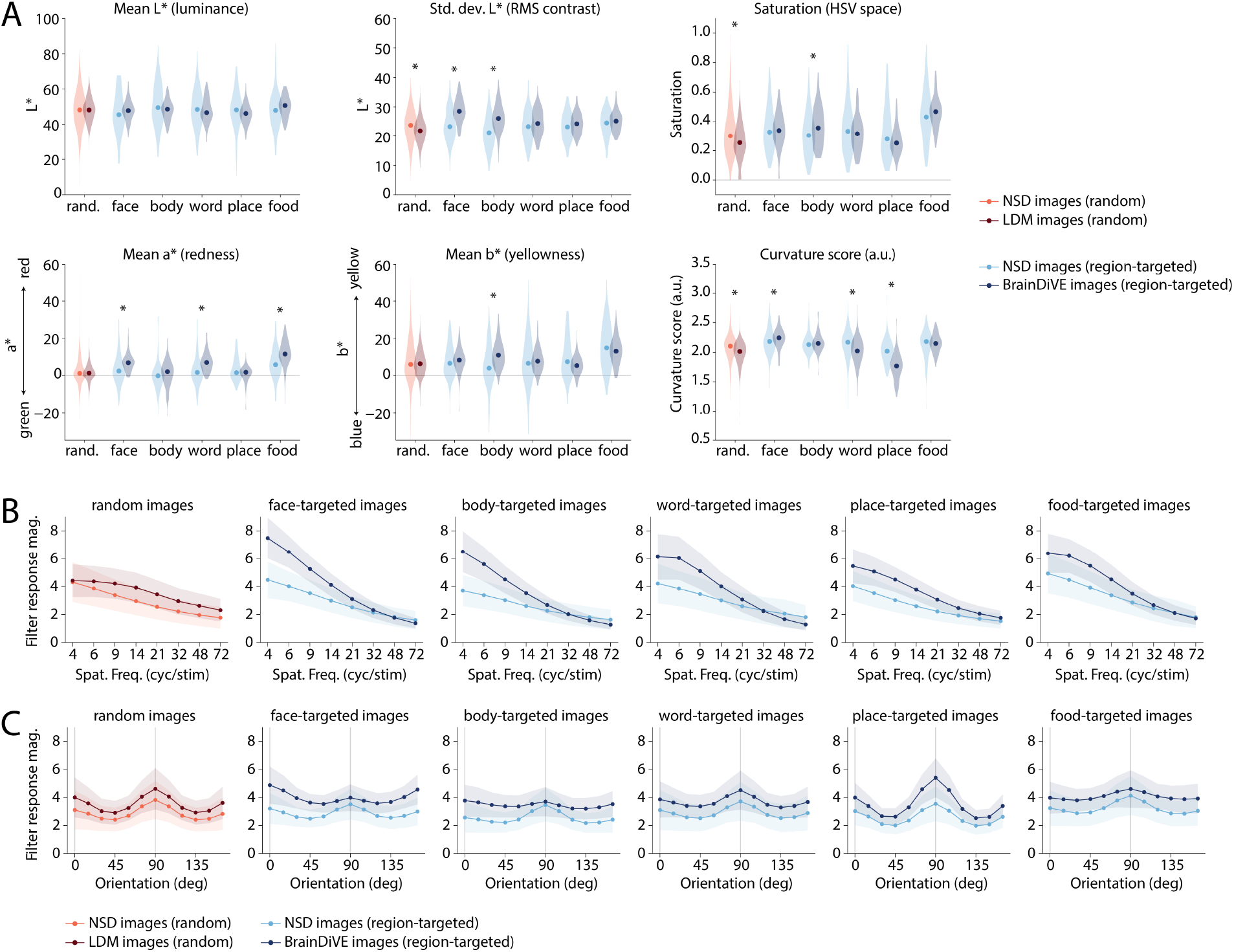
Low-level and mid-level image statistics of ROI-targeted images. **(A)** For each stimulus image, we quantified the luminance (mean L*), RMS contrast (std. dev. L*), color saturation, color hue along the greenred (a*) and the blue-yellow (b*) opponency axes, and estimated edge curvature (see *Methods* for details on quantification). As a baseline comparison, we also computed statistics for a group of 1000 randomly-selected NSD images (originally taken from the COCO dataset; Lin et al., 2014) and for a group of 1000 random synthetic images generated using Stable Diffusion 2.1 base with no brain guidance (see *Methods* for details). Violin plots show the distributions of these statistics across image groups. Violins in red shades indicate the natural and synthetic baseline images (rand. = random); blue shades indicate 100 NSD images and 100 BrainDiVE images targeted to each ROI; dots indicate medians. Asterisks indicate a significant difference between the distributions for natural versus model-generated images (Mann-Whitney *U*-test, *p <* 0.01). **(B-C)** The orientation and spatial frequency content of images were quantified using a Gabor filter bank, see *Methods*. **(B)** Shows the average Gabor response magnitude as a function of spatial frequency, averaged across orientation. **(C)** Shows magnitude as a function of orientation, averaged across spatial frequency. 0°= vertical, 90°= horizontal. Dots and shaded error bars indicate the mean and standard deviation across images in each group.

These analyses revealed region-dependent differences between BrainDiVE and NSD images, consistent with variation in low- and mid-level feature selectivity across regions. Considering low-level statistics (Figure 4A), luminance did not differ between NSD and BrainDiVE images. However, BrainDiVE images targeted to face- and body-selective regions showed higher RMS contrast than the corresponding NSD images. This difference was not observed for the random LDM-generated images relative to the random NSD images, suggesting that the effect is not an artifact of the LDM synthesis procedure, but may reflect sensitivity of these regions to contrast. No differences in RMS contrast were observed between NSD and BrainDiVE images targeted to word-, place-, or food-selective ROIs.

For color statistics, BrainDiVE images targeting face-, word-, and food-selective regions were shifted toward higher red values (a* in CIEL* a* b* space) relative to NSD images. BrainDiVE images targeting body-selective regions also showed higher yellow values (b*) and greater saturation relative to NSD images. These color differences were not evident when comparing random LDM-generated versus random NSD images. We also observed that for both NSD and BrainDiVE, images targeting food-selective ROIs had higher values of a*, b*, and saturation values than images targeting other ROIs. Together, these results suggest that warm, saturated colors contribute to responses in face-, body-, word-, and food-selective ROIs, partially consistent with prior work (Duyck et al., 2021; Nestor & Tarr, 2008; Pennock et al., 2023).

With respect to spectral statistics, we observed ROI-dependent differences between BrainDiVE and NSD images. Across all categories, BrainDiVE images showed an increase in power at low spatial frequencies compared to NSD images (Figure 4B). In contrast to BrainDiVE images, random LDM-generated images tended to exhibit greater power at high spatial frequencies, consistent with prior observations of elevated high-frequency detail in latent diffusion models (Corvi et al., 2023; Lee & Chang, 2022; Rombach et al., 2022). This indicates that the low-frequency shift in BrainDiVE images is not solely driven by the LDM synthesis procedure but may reflect neural tuning to spatial frequency. Moreover, the extent of the bias toward low frequencies in the BrainDiVE images differed across regions, with face- and body-targeted BrainDiVE images showing a stronger bias toward low spatial frequencies, and place-targeted BrainDiVE images showing a weaker bias. This points towards differential sensitivity to low spatial frequencies across regions, consistent with prior work (Henderson et al., 2023; Rajimehr et al., 2011).

We next examined the orientation distribution across images (Figure 4C). NSD images targeting all regions, as well as random NSD images, exhibited a cardinal bias, with greater magnitude at vertical and horizontal orientation relative to diagonal, as expected for natural images (Girshick et al., 2011; Henderson & Serences, 2021). A similar cardinal bias was observed for random LDM-generated images.

In contrast, the orientation distribution for BrainDiVE images varied across ROIs. For place-selective ROIs, BrainDiVE images exhibited increased power at horizontal orientation (90°) relative to diagonal, leading to a more peaked distribution compared to place-targeted NSD images. For face-, body-, and food-selective ROIs, BrainDiVE images exhibited a flatter orientation distribution compared to NSD images, with increased power at vertical and diagonal orientation, particularly at low spatial frequencies (Supplementary Figure 2).

Finally, in terms of curvature statistics, BrainDiVE images targeting face-selective regions exhibited higher edge curvature than NSD images, whereas BrainDiVE images targeting word- and place-selective regions exhibited lower edge curvature. For the random image groups, LDM-generated images also exhibited lower edge curvature than NSD images, suggesting that the diffusion process may account for part of the difference between BrainDiVE and NSD images targeting word- and place-selective ROIs. However, the decrease in curvature from NSD to BrainDiVE images for place-selective ROIs was larger than that observed for random images, suggesting neural selectivity contributed to this effect. This pattern is consistent with face- and place-selective regions being biased toward curved and rectilinear contours, respectively (Nasr et al., 2014; Ponce et al., 2017; Yue et al., 2020a). The implications of these findings are discussed further in the *Discussion*.

### Differentiating face-selective regions

In addition to differentiating broad category-selective networks, BrainDiVE-generated images reveal differences between individual face-selective ROIs, namely FFA and OFA. While FFA-targeted and OFA-targeted images both contained faces or face-like structures, FFA-targeted images more often depicted complete, realistic faces, whereas OFA-targeted images typically included face-like elements that were less photorealistic (Figure 5A). We quantified this difference using both image statistics and behavioral data (Figure 5B-C; see *Methods*). Analysis of mean curvature, luminance, color, and spectral statistics showed that FFA-targeted BrainDiVE images had higher RMS contrast (SD of L*) than OFA-targeted images, but no other statistics differed between groups (Figure 5B, Supplementary Figure 2). This suggests that the differences between these regions primarily reflect high-level, semantic content rather than low- or mid-level image properties. Consistent with this, our behavioral study showed that FFA-targeted images were rated as more photorealistic and as containing more human faces, whereas OFA-targeted images were rated as containing more animal-like faces, abstract shapes, text, and non-face objects (Figure 5C). While similar differences were present in the top-activating NSD images for FFA and OFA, the magnitude of these effects was larger for BrainDiVE images, suggesting that BrainDiVE better captures the image properties that differentiate these regions.

**Figure 5.**
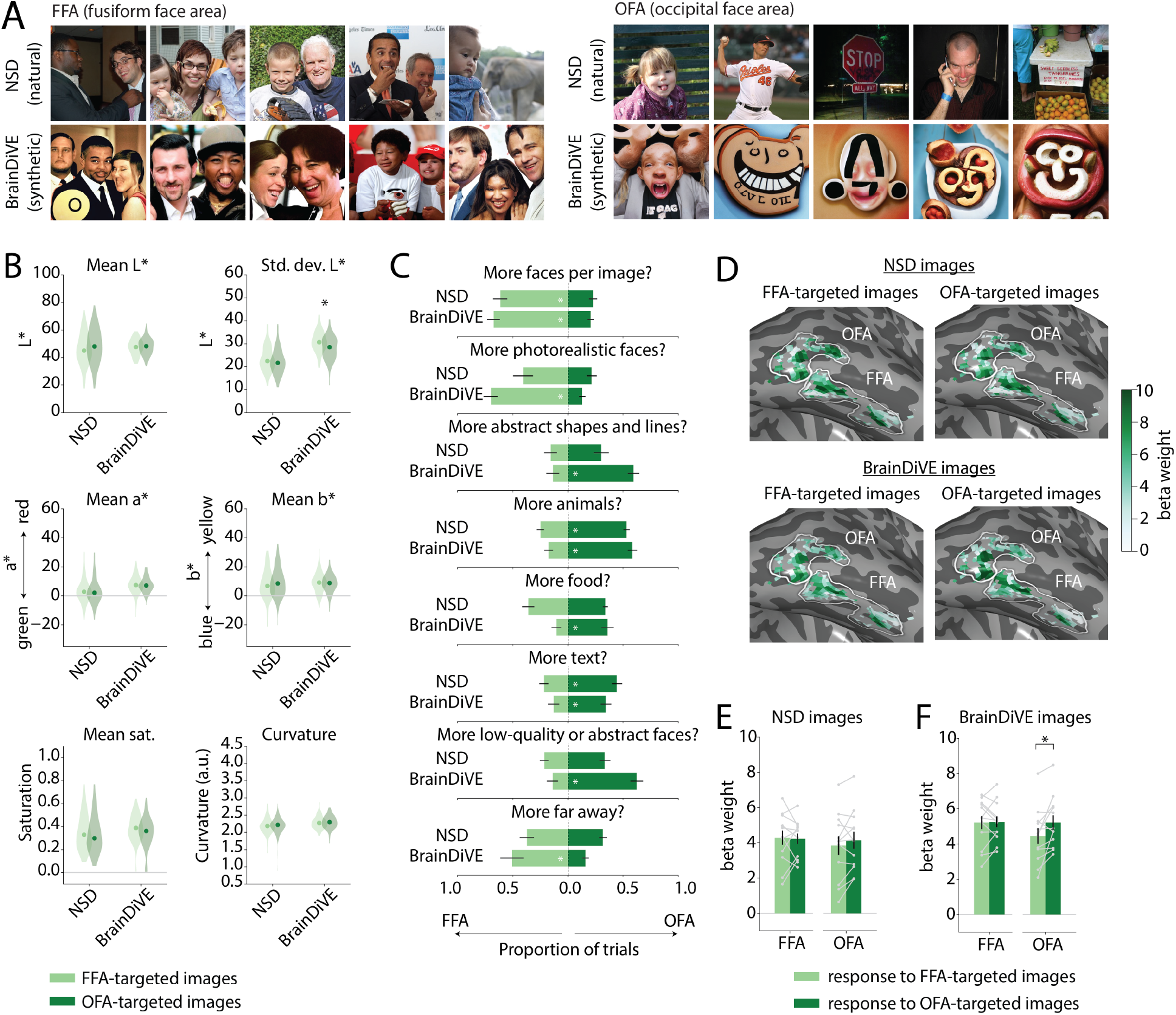
Model-generated images differentiate regions within the face-processing network. **(A)** Separate image sets were generated to target the FFA (left) and the OFA (right); images shown are for NSD participant S2. In each panel, the top row shows the five highest activating natural images (NSD stimuli viewed by S2), and the bottom row shows the five highest activating BrainDiVE images. **(B)** Image statistics for FFA- and OFA-targeted images. Violin plots show distributions across 100 NSD or BrainDiVE images targeted per region; dots indicate medians (see *Methods* and Figure 4 caption for more details on metrics); asterisks significant differences between FFA and OFA distributions (Mann-Whitney *U*-test, *p <* 0.01). **(C)** Behavioral results in which participants (n=10) compared groups of FFA- and OFA-targeted images from NSD or BrainDiVE and decided which group rated more highly along various perceptual and semantic dimensions (see *Methods*). Participants could also select “same” for each dimension; these responses are not shown. Bar lengths and error bars indicate mean and SEM across participants; asterisks indicate significant differences in the proportion of trials on which FFA-versus OFA-targeted images were chosen (paired *t*-test with permutation; all *p <* 0.01, see *Methods*). **(D)** Voxel responses (beta weights) from an example participant (S03) shown on an inflated cortical surface. Panels show responses to FFA- or OFA-targeted images (NSD and BrainDiVE). Voxels are thresholded based on face selectivity in an independent localizer (*t >* 2). White outlines indicate functionally-defined ROIs (see *Methods*). **(E-F)** Mean responses (beta weights) of FFA and OFA to FFA- and OFA-targeted images across participants (n=12), shown separately for **(E)** NSD and **(F)** BrainDiVE runs. Bar heights and error bars indicate mean and SEM across participants; gray points and lines show individual participant data; asterisks indicate significant differences between responses to FFA- and OFA-targeted images (paired *t*-test with permutation; all *p <* 0.05; see main text for test values). See Supplementary Figure 3 for results for all ROIs. All photographs, including those depicting people, are from the Microsoft COCO dataset (Lin et al., 2014) and are used in accordance with the COCO dataset’s Creative Commons Attribution 4.0 license.

Returning to our fMRI results, we evaluated the consistency of the FFA/OFA difference across participants by comparing responses in FFA and OFA (as defined in our new participants) evoked by images targeting these regions in NSD participants. Our results demonstrate the cross-participant generalizability of the FFA/OFA difference (Figure 5D-F). As expected given the face-like appearance of both FFA- and OFA-targeted images, we observed positive average responses (*β* weights) to both FFA-targeted and OFA-targeted images in FFA and OFA, for both top-activating NSD images (Figure 5E) and BrainDiVE images (Figure 5F). When considering BrainDiVE images only, OFA was more responsive to OFA-targeted than FFA-targeted images, whereas FFA was equally responsive to both image sets. A two-way repeated measures ANOVA revealed a significant interaction between ROI and Image Type (FFA-vs. OFA-targeted images), with no significant main effects (two-way repeated measures ANOVA;BrainDiVE;ROI: *F(* 1, 11) = 0.94, *p* = 0.364;Image Type: *F* (1, 11) = 2.66, *p* = 0.134;ROI x Image Type: *F* (1, 11) = 25.82, *p <* 0.0001; *p*-values obtained using a permutation test; see *Methods*). Focused *t*-tests showed that responses to OFA-targeted BrainDiVE images were significantly greater than those to FFA-targeted images in OFA (two-tailed paired *t*-test with permutation: *t* (11) = −2.78, *p* = 0.015), but not in FFA (*t* (11) = −0.16, *p* = 0.857). This result indicates that differences between FFA- and OFA-targeted BrainDiVE images correspond to a functional difference between these ROIs: OFA-targeted images, with a larger proportion of abstract, non-realistic face features, elicited larger responses in OFA than FFA-targeted images. In contrast, responses to NSD images did not differ between FFA- and OFA-targeted images (two-way repeated measures ANOVA;NSD;ROI: *F* (1, 11) = 0.33, *p* = 0.587;Image Type: *F* (1, 11) = 0.24, *p* = 0.648;ROI x Image Type: *F* (1, 11) = 2.58, *p* = 0.134; *p*-values obtained using permutation test; see *Methods*). This suggests that BrainDiVE isolates functional differences in FFA and OFA that are not captured by simply selecting the highest-activating natural images for each region.

### Differentiating subregions of occipital place area

At an even finer level of granularity, our previous results (Luo et al., 2023) suggested that BrainDiVE can differentiate the functional properties of two clusters of voxels within the scene-selective region OPA. Using voxel clusters functionally identified in NSD participants (see *Methods*), we defined a posterior cluster (“OPA1”) that appeared to be selective for indoor, manmade scenes and an anterior cluster (“OPA2”) selective for outdoor, natural scenes (Figure 6). Image statistics for the BrainDiVE image sets associated with these two clusters revealed that OPA1-targeted images were warmer in color (higher a* and b*) and higher in RMS contrast (SD of L*) than OPA2-targeted images, while OPA-2 targeted images were brighter (higher L*) and had higher estimated curvature (Figure 6C). A behavioral study found significant perceptual differences between these clusters. OPA1-targeted images were rated as more indoor and containing more angular, geometric features, while OPA2-targeted images were rated as more natural and further away (i.e., larger scale;Figure 6D).

**Figure 6.**
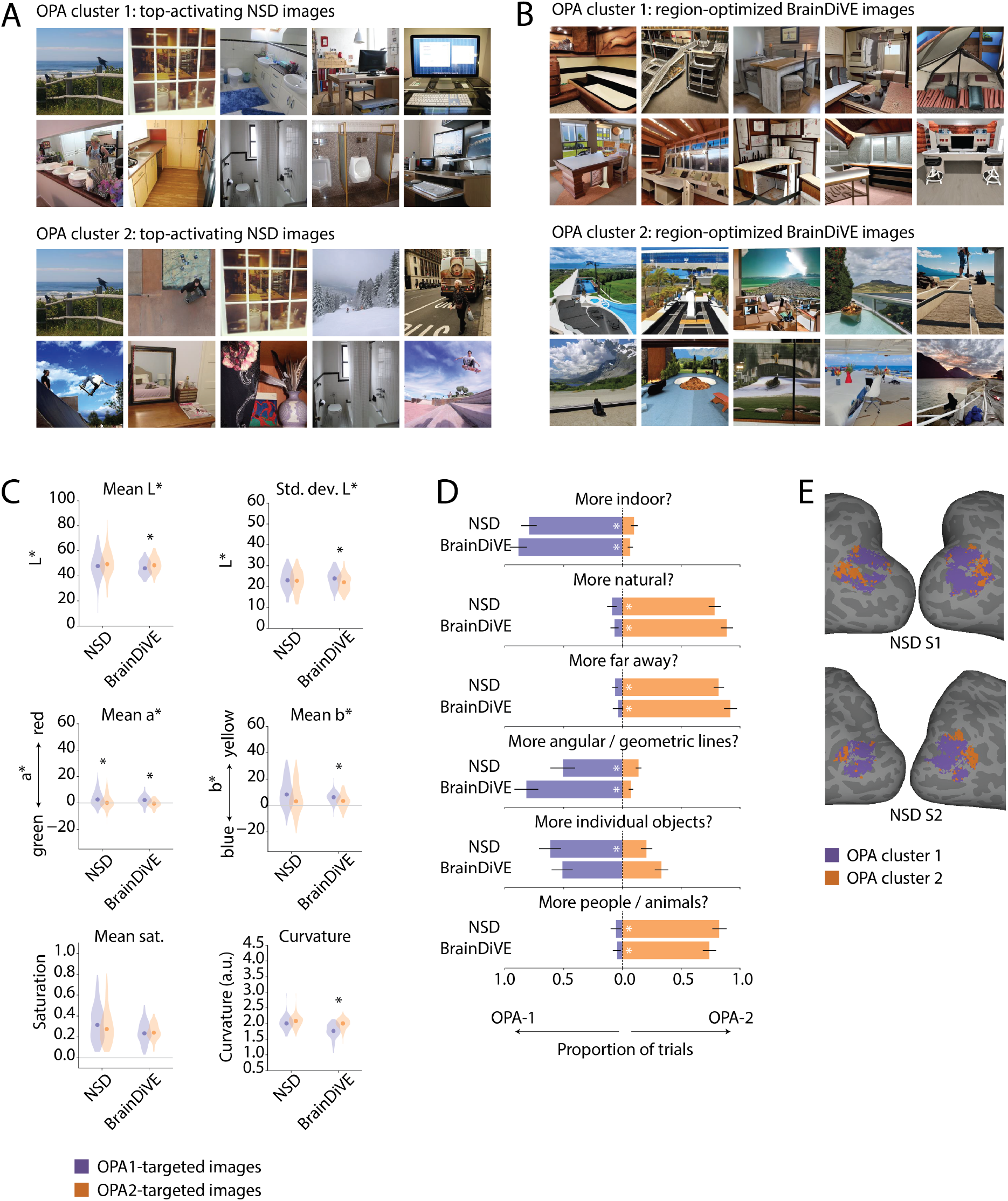
Subregions of the occipital place area (OPA) are responsive to different scene properties. Subregions (OPA1 and OPA2) were identified in NSD participants using vMF clustering, and image sets were created to target OPA1 and OPA2 separately (see *Methods*). Example images shown here were selected based on NSD participant S2. **(A)** Top-10 activating natural NSD images shown to S2, for each OPA subregion. **(B)** Top-10 images for each subregion, generated using BrainDiVE. **(C)** Quantifying image statistics across OPA1- and OPA2-targeted images. Violin plots show the distribution of each visual property across 100 NSD or BrainDiVE images targeted to each subregion; dot shows the median (for details on metrics see *Methods* and Figure 4). Asterisks denote a significant difference between OPA1- and OPA2-targeted distributions (Mann-Whitney *U*-test, *p <* 0.01). **(D)** Behavioral results in which participants (n=10) compared groups of OPA1- and OPA2-targeted images from NSD or BrainDiVE and decided which group of images rated more highly along various perceptual and semantic dimensions (see *Methods*). Bar lengths and error bars indicate mean and SEM across participants; asterisks indicate significant differences in the proportion of trials on which OPA1-versus OPA2-targeted images were chosen (paired *t*-test with permutation; all *p <* 0.01, see *Methods*). **(E)** Locations of OPA1 and OPA2 clusters in two NSD participants on an inflated cortical surface (Luo et al., 2023). All photographs, including those depicting people, are from the Microsoft COCO dataset (Lin et al., 2014) and are used in accordance with the COCO dataset’s Creative Commons Attribution 4.0 license.

To assess whether OPA1 and OPA2 could be identified in our current participants, we examined the anatomical organization of responses to OPA1- and OPA2-targeted images (Figure 7). First, for NSD and BrainDiVE images separately, we computed the contrast for OPA2-versus OPA1-targeted images. Figure 7A shows these *t*-statistics on an inflated cortical surface for one participant. Consistent with the locations of these clusters in NSD participants, we observed a spatial gradient: posterior OPA showed stronger responses to OPA1-targeted images, whereas anterior OPA showed stronger responses to OPA2-targeted images. To quantify this functional gradient, we assigned each surface space vertex a coordinate based on its distance from posterior and medial anatomical landmarks, yielding position indices for each vertex along posterior-to-anterior and medial-to-lateral anatomical gradients within OPA (see *Methods* for details). Visualizing this 2D coordinate space reveals the same spatial organization evident in the cortical surface map (Figure 7B). We next performed k-means clustering on the *t*-statistic values and computed cluster medians (Figure 7C). Visualizing cluster medians across participants and hemispheres (Figure 7D) revealed spatial segregation that was consistent across participants. For cluster medians derived from NSD images (Figure 7D, left), voxels more responsive to OPA1-targeted images (“Cluster 1”) were located more posteriorly, whereas voxels more responsive to OPA2-targeted images (“Cluster 2”) were located more anteriorly. In contrast, cluster medians showed less separation along the medial-to-lateral axis. When the same clustering was applied to responses evoked by BrainDiVE images, the posterior-to-anterior organization was less apparent (Figure 7D, right). Although some participants showed a similar pattern for OPA1- and OPA2-targeted BrainDiVE images as observed with NSD, it was less consistent across participants.

**Figure 7.**
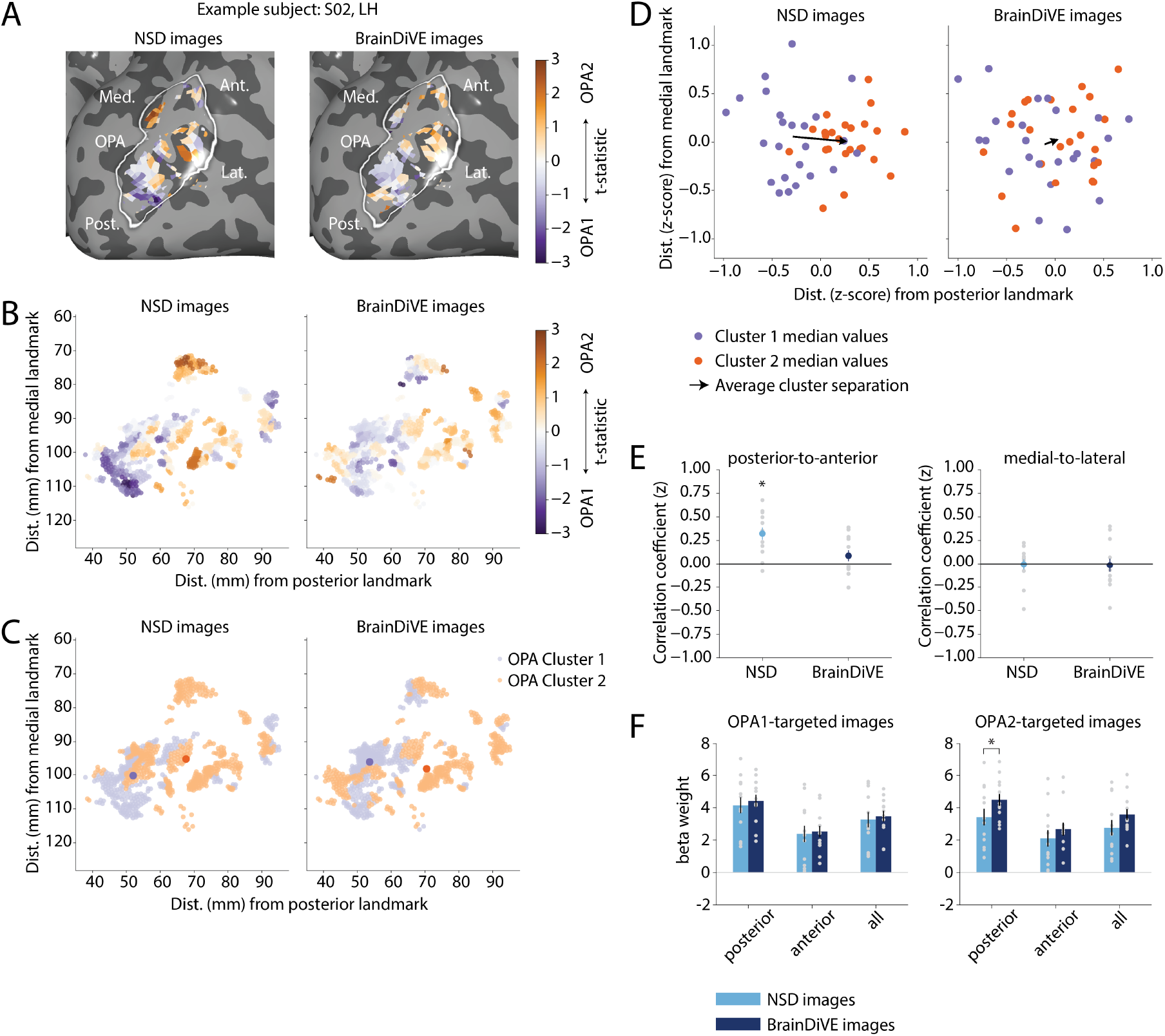
Images targeted to subregions of the occipital place area (OPA) yield spatially distinct response profiles. OPA subregions (OPA1 and OPA2) were identified in NSD participants using vMF clustering (see *Methods*), and separate sets of images were generated to target each subregion (Figure 6). **(A)** Differences in responses to OPA2-versus OPA1-targeted images (*t*-statistic), for one participant (S02), on an inflated left cortical surface (surface is left-right mirrored). OPA voxels were selected in an independent functional localizer (localizer *t >* 2). Left: *t*-statistics for NSD images; right: *t*-statistics for BrainDiVE images. **(B)** Vertex data from (A) plotted as a function of anatomical position along posterior-to-anterior and medial-to-lateral axes (see *Methods*). Each dot denotes a surface vertex. **(C)** Results of k-means clustering analysis (k=2) on OPA *t*-statistics from (B), showing spatial separation of clusters. Colors indicate different clusters; larger dots denote cluster medians. **(D)** Average cluster separation across participants and hemispheres. Cluster medians were computed after z-scoring vertex positions to account for differences in the absolute positions of OPA across participants. Each dot represents one hemisphere in one participant; colors indicate clusters. Black arrows show vectors between the mean cluster medians (i.e., average cluster separation). **(E)** Correlations between the OPA2 *>* OPA1 *t*-statistic for individual vertices and the anatomical position along the posterior-to-anterior (left) and medial-to-lateral (right) axis. Gray dots show correlation coefficients (Fisher z-transformed) for single participants; colored dots and error bars show mean and SEM across participants. Asterisks indicate correlations significantly different from zero (one-sample *t*-test with permutation, *p <* 0.01, see *Methods*). See Supplementary Figure 5 for *t-*statistic versus vertex position scatter plots. **(F)** Responses to OPA1- and OPA2-targeted images for posterior OPA, anterior OPA, and all vertices combined (all). Posterior and anterior regions were defined using a median split on anatomical distance values; see *Methods* for details. Gray dots show individual participants; bar heights and error bars indicate mean and SEM across participants. Asterisks indicate significant differences between NSD and BrainDiVE responses (paired *t*-test with permutation, *p<*0.01, see *Methods*). See Supplementary Figure 4 for additional comparisons.

To quantify these effects, we computed the correlation across vertices between *t*-statistics for the contrast of OPA2-versus OPA1-targeted images, and each vertex’s position along the anatomical axes. For NSD images, this revealed a consistent positive relationship with posterior-to-anterior position (Figure 7E; Supplementary Figure 5), indicating relatively greater responses to OPA2-targeted images in more anterior vertices. The mean of the correlation coefficients for individual participants was significantly greater than zero (one-sample *t*-test: *t* (11) = 4.78, *p* = 0.003; two-tailed *p*-value computed using a permutation test; see *Methods*), indicating that this anatomical gradient was consistent across participants. For BrainDiVE images, a similar positive relationship was evident, but the mean correlation coefficient was not significantly different from zero (*t* (11) = 1.47, *p* = 0.171). No significant relationship was observed between vertex *t*-statistics and the medial-to-lateral position of each vertex for either image set (one-sample *t*-tests;NSD: *t* (11) = −0.12, *p* = 0.895;BrainDiVE: *t* (11) = −0.20, *p* = 0.827; two-tailed *p*-values computed using a permutation test; see *Methods*).

These results indicate a difference between responses evoked by natural and model-generated images: NSD images reliably elicited the expected posterior-to-anterior organization within OPA, whereas BrainDiVE images exhibited greater variability. To better understand this effect, we examined responses within OPA to OPA1- and OPA2-targeted images separately. Given that the strongest functional organization was along the posterior-to-anterior axis, we performed a median split along this axis, dividing OPA vertices in each participant and hemisphere into posterior and anterior groups. Because these groups were defined purely anatomically, they enabled an unbiased comparison of responses to NSD and BrainDiVE images (Figure 7F and Supplementary Figure 4).

This analysis revealed an off-target effect for OPA2-targeted BrainDiVE images, in that these images elicited stronger responses in posterior OPA than OPA2-targeted NSD images (BrainDiVE *t* vs. NSD *t*; t(11) = 3.53, *p* = 0.008; two-tailed *p*-values computed using a permutation test; see *Methods*). In contrast, anterior OPA, where OPA2-targeted images were expected to drive responses, did not show a significant difference between NSD and BrainDiVE images (*t* (11) = 2.06, *p* = 0.064). No differences were observed for OPA1-targeted images for either posterior or anterior vertices.

When directly contrasting responses to OPA1-versus OPA2-targeted images (Supplementary Figure 4), we observed that posterior OPA was significantly more responsive to OPA1-versus OPA2-targeted NSD images (*t* (11) = 4.64, *p* = 0.000). In contrast, for BrainDiVE images, neither posterior nor anterior OPA differentiated between OPA1 and OPA2-targeted images, reflecting the off-target posterior response to OPA2-targeted BrainDiVE images. These patterns may explain the lack of consistent anatomical organization when comparing responses to OPA1- and OPA2-targeted BrainDiVE images. We consider potential explanations for this effect in our *Discussion*.

## Discussion

Subregions of higher visual cortex reliably respond to specific image categories, but it remains challenging to identify the perceptual and semantic properties that drive these responses. Computational stimulus optimization provides a tool for disentangling these tuning dimensions by visualizing images associated with maximal responses. Here, we developed and validated BrainDiVE, a stimulus optimization method for higher visual cortex, and used it to characterize the key image properties underlying category-selective responses. For face-, body-, word-, and place-selective regions, BrainDiVE-generated images elicited higher category selectivity than natural images targeted to the same regions, indicating that our method successfully isolates the image space dimensions that drive each region. We further show that such feature axes can be derived not only for broad category-selective networks but also for pairs of ROIs responsive to the same category (OFA, FFA). As discussed below, these findings inform and enhance our understanding of the functional organization in higher visual cortex.

### Differences between natural and model-generated images

Comparing BrainDiVE and NSD images targeting the same cortical regions provides a measure of the stimulus dimensions most strongly associated with neural responses. Across several category-selective regions, BrainDiVE images appear to exaggerate characteristics of the represented category. For example, word-targeted BrainDiVE images contain more text-like elements than their NSD counterparts, and place-targeted BrainDiVE images contain more prominent structural elements and fewer objects than their NSD counterparts. In addition to these semantic differences, BrainDiVE images also exaggerate specific low- and mid-level features relative to NSD images. For example, face-targeted BrainDiVE images contain higher curvature relative to their NSD counterparts, whereas place- and word-targeted BrainDiVE images contain lower curvature relative to their NSD counterparts. These findings are consistent with prior evidence on the large-scale organization of curvature within higher visual cortex, with face-selective regions preferring curvilinear edges and scene-selective regions preferring rectilinear edges (Nasr et al., 2014; Ponce et al., 2017; Yue et al., 2020a, 2020b). More broadly, our results provide converging evidence for the role of curvature and rectilinearity in shaping cortical selectivity, showing that images that exaggerate these features elicit stronger responses.

In addition to curvature, our results indicate differential sensitivity to spectral statistics and color across category-selective ROIs. Considering spectral statistics, place-targeted BrainDiVE images showed greater power at horizontal orientations than NSD images. In contrast, BrainDiVE images targeting other regions showed flatter distributions and/or greater power at vertical orientations. Prior work demonstrated selectivity for cardinal orientations (vertical and horizontal) in scene-selective ROIs, consistent with our results, but has not reported an asymmetry between horizontal and vertical orientations (Henderson et al., 2023; Nasr & Tootell, 2012). Our results suggest that scene-selective regions may be particularly sensitive to horizontally-oriented scene elements (e.g., the horizon), consistent with evidence that horizontal orientations are overrepresented in natural scenes and that orientation perception exhibits vertical/horizontal asymmetries (Essock et al., 2003). Additionally, BrainDiVE images targeted to all ROIs exhibited greater power at low spatial frequencies than NSD images, an effect which was not observed in randomly generated control images. This low-frequency bias was largest for face- and body-targeted BrainDiVE images and smallest for place-targeted BrainDiVE images, suggesting differential sensitivity to low spatial frequencies across different category-selective networks. Consistent with this, prior work has shown that scene-selective regions such as PPA and RSC are biased to higher spatial frequencies (Berman et al., 2017; Henderson et al., 2023; Kauffmann et al., 2015; Rajimehr et al., 2011), whereas face-selective regions are selective for lower spatial frequencies (Henderson et al., 2023).

With respect to color, face-, word-, and food-targeted BrainDiVE images showed higher red values than NSD images. In contrast, body-targeted BrainDiVE images showed higher yellow values and greater saturation. This finding in food-selective regions is consistent with evidence that these regions overlap with cortical populations selective for saturated color (Lafer-Sousa et al., 2016; Pennock et al., 2023; Rosenthal et al., 2018). Our results also suggest a role for color in shaping selectivity in face-, body-, and word-selective regions. It is possible that the precise distribution of colors identified in face-targeted and body-targeted BrainDiVE images could reflect biases in the skin color of people depicted in our model training data (COCO, LAION-5B; Zhao et al., 2021). Prior work has demonstrated warm-biased color tuning in macaque face-selective patches (Duyck et al., 2021), but the overall role of color in the face-, body-, and word-processing networks remains poorly understood, highlighting a direction for future work.

### Scene-selective regions

The patterns of differential responses to BrainDiVE versus NSD images varied across category-selective networks. Face-, body-, and word-selective regions all showed a significant increase in response from NSD to BrainDiVE for “on-target” images (i.e., images targeting the corresponding network). In contrast, place-selective regions did not show a significant response difference from NSD to BrainDiVE for on-target images. Instead, both OPA and PPA exhibited reduced response to “off-target” BrainDiVE images (i.e., images targeting other networks). This was reflected in increased “place selectivity” (i.e., *t*-statistic in Figure 3D) from NSD to BrainDiVE. This pattern may reflect properties of the stimulus set. In particular, NSD images are natural scenes that typically contain some background and scene structure, such that the highest activating NSD images may already be approaching the optimal stimulus for scene-selective cortex, limiting further response increases. In contrast, for regions selective for faces, bodies, and words, even the most effective NSD images often contain scene-related content, leading to off-target activation in PPA and OPA. BrainDiVE appears to suppress this scene-related content when targeting non-scene ROIs, generating images that more effectively isolate category-relevant features. Overall, these results highlight the region specificity of BrainDiVE: images targeting a given region evoke spatially-specific activation patterns with reduced off-target responses relative to NSD images (see also Figure 2). This specificity suggests that the method distills the components of natural images that most effectively drive distinct category-selective networks.

In the occipital place area (OPA), we found evidence for a functional distinction between the posterior and anterior portions, which was reliably detectable when using NSD but not BrainDiVE images. In terms of image properties, this distinction indicated greater selectivity for indoor, angular scenes in the more posterior cluster (which we termed OPA1), and greater selectivity for outdoor, natural, distant scenes in the more anterior cluster (OPA2). This finding is consistent with some results from prior work, which suggested a gradient in OPA reflecting an axis from closed-to-open scenes, along a similar anatomical direction (Lescroart & Gallant, 2019). While we did not obtain perceptual measurements of scene openness, visual inspection of the OPA1-targeted and OPA2-targeted images (Figure 6) suggests openness could be another image dimension driving this organization. Other work has similarly suggested a gradient within OPA related to scene scale (Peer et al., 2019). In our results, this anatomical organization of responses to OPA1-targeted and OPA2-targeted NSD images was consistent across participants (Figure 7), even though the stimulus images were selected based on completely independent participants in NSD. Thus, our result provides novel evidence strengthening the claim of intra-region organization within the OPA and its generalizability across subjects. However, when using BrainDiVE images that were separately targeted to OPA1 and OPA2, we observed less consistency in organization, with OPA2-targeted images giving “off-target” activation in posterior OPA (Figure 7F). This suggests the OPA1-targeted and OPA2-targeted BrainDiVE images were not sufficiently specific to differentially target these subregions. Because OPA1 and OPA2 responses are likely highly correlated in NSD, this result indicates new computational strategies may be needed to precisely disentangle the tuning of such correlated regions.

### Food-selective regions

Comparing NSD versus BrainDiVE images, we did not observe an increased response of food-selective regions to food-targeted images, nor an increase in the *t*-statistic contrasting food-targeted versus other images. However, responses to word-targeted images in food-selective regions did significantly increase from NSD to BrainDiVE. This effect likely reflects functional overlap between food- and word-selective ROIs in the NSD analysis. The ROI masks used during image generation from NSD were relatively large (localizer *t*-statistic *>* 2), and did not exclude overlap across categories, such that voxels with both food- and word-selectivity may have contributed to the generation of word-targeted BrainDiVE images (note that in our new experimental data, ROI definitions did exclude category overlap). As a result, the word-targeted images may contain some food-related features, consistent with their increased redness and some aspects of their apparent content (Figure 1). Notably, this effect was not reciprocal: food-targeted BrainDiVE images did not elicit increased responses in word-selective regions. This asymmetry may suggest broader tuning of neural populations in food-selective regions and more selective tuning in word-selective regions. Given that food-selective ROIs were defined in NSD, which includes relatively few word-related images, further work is needed to disentangle the relationship between food- and word-selectivity in ventral visual cortex. Finally, although responses to food-targeted images did not differ between BrainDiVE and NSD, both image sets elicited significant food selectivity across participants, comparable in magnitude to that observed in other category-selective ROIs (positive *t*-statistics in Figure 3E). These results provide converging evidence for the reliability of food-selective responses in these recently identified regions.

### Face-selective regions

BrainDiVE images differentiated responses in the occipital face area (OFA) and the fusiform face area (FFA): OFA showed greater response to OFA-targeted than FFA-targeted BrainDiVE images, whereas FFA responded similarly to both. This difference was not detectable with NSD images, highlighting the utility of BrainDiVE for distinguishing responses in similarly tuned ROIs. OFA-targeted BrainDiVE images were more abstract than FFA-targeted images, often containing cartoon-like features resembling face parts. This pattern is consistent with accounts in which OFA encodes local face features and FFA encodes more global face configurations (Gauthier et al., 2000; Grill-Spector et al., 2017; Liu et al., 2010; Pitcher et al., 2011). However, prior work suggests that OFA responds similarly to face parts regardless of configuration, whereas FFA is more sensitive to specific configurations (Liu et al., 2010), which would predict comparable OFA responses to both FFA-targeted and OFA-targeted images. Instead, OFA preferentially responded to OFA-targeted BrainDiVE images, whereas FFA showed increased responses to BrainDiVE images regardless of target (Supplementary Figure 3).

One explanation for this pattern of results is that OFA-targeted BrainDiVE images differ from the scrambled faces used in prior studies, as they are not explicitly constructed to disrupt spatial configuration. Because BrainDiVE relies on a pretrained latent diffusion model with a strong prior over natural images, these images may retain configural information despite their abstract appearance. More generally, these results suggest that OFA is selective for mid-to high-level features associated with faces that are preferentially amplified in OFA-targeted images. The lack of differentiation in FFA may reflect increased invariance at later stages of the face-processing network, with reduced sensitivity to these OFA-specific features.

### Prior and future work

Recent work has explored the use of generative models for neural stimulus optimization. Approaches such as NeuroGen use generative adversarial networks (GANs) with encoding models to generate region-optimized stimuli (Gu et al., 2022; Ratan Murty et al., 2021), while more recent approaches combine diffusion models with transformer-based encoding models to probe fine-grained semantic selectivity (Hwang et al., 2025). BrainACTIV instead manipulates reference images to maximize or minimize activation in target brain regions (Garcia Cerdas et al., 2025). Notably, both NeuroGen and BrainACTIV report differences between OFA and FFA that are broadly consistent with our findings, including text-like or animal-like features in OFA-targeted images. Related work has shown that model-derived stimuli can be used to parametrically control neural responses along encoding axes, revealing differences in feature tuning and model-brain alignment not captured by standard methods (Prince et al., 2025). This approach provides a complementary framework for probing neural tuning by systematically manipulating images along targeted feature dimensions, and, as in our study, can generate testable predictions for new fMRI participants.

Our results extend prior work by demonstrating that model-generated images can selectively and more strongly activate target cortical populations in new human participants. This contrasts with findings from NeuroGen, which reported similar activation for natural and model-optimized stimuli in higher visual cortex (Gu et al., 2023). This discrepancy may reflect methodological differences, including the use of GANs rather than diffusion models and an fMRI design that included longer blocks and greater spacing between images. Our design, with shorter blocks and denser image onsets, may have increased sensitivity to detect subtle differences between image types. This is consistent with NeuroGen’s report of advantages for model-generated versus natural images in some ROIs (e.g., anterior temporal lobe/aTL-faces, and fusiform body area/FBA1; Gu et al., 2023). These specific ROIs may have larger effect sizes or be more aligned with biases in NeuroGen’s image priors. In related work, other stimulus optimization frameworks have demonstrated in mice and non-human primates that optimized images can drive neural responses beyond those evoked by natural images (Bashivan et al., 2019; Cowley et al., 2026; Ponce et al., 2019; Walker et al., 2019). Our results extend these findings to non-invasive measurements in humans, demonstrating effective stimulus optimization in semantically selective neural populations within higher visual cortex.

The ability to elicit spatially specific activation patterns in target neural populations using non-invasive measurements provides a new avenue for controlling and probing visual cortical representations. Future variants of our approach should be capable of targeting increasingly precise neural subpopulations or modulating activity along continuous visuo-semantic dimensions, enabling fine-grained control over neural responses (Prince et al., 2025). Combined with advances in real-time fMRI (Leeds & Tarr, 2016; Sitaram et al., 2017; Sulzer et al., 2013), our framework opens the possibility of a fully closed-loop stimulus optimization framework for participant-specific control of brain activity. More broadly, BrainDiVE establishes a data-driven framework for understanding –and ultimately controlling –cortical representations. In this spirit, our work moves beyond piecemeal probing of neural systems toward more systematic, model-based interrogation of cortical function (Newell, 1998).

## Methods

### Human participants

Participants were adults recruited from the Carnegie Mellon community and had normal or corrected-to-normal vision. Data was collected from 16 participants in total;12 of the 16 were included in our final dataset (average age of 12 final participants: 23.92 ±5.12;mean ±SD). Three participants were excluded from our analyses because of excessive head motion during functional runs and a fourth participant was excluded due to mild tissue heating that occurred during the scan session. Each participant completed one 90 minute MRI session and was compensated $30 per hour. All participants provided written informed consent, and the protocol was approved by the Institutional Review Board at Carnegie Mellon University.

### Acquisition of fMRI data

MRI data was collected at the CMU-Pitt Brain Imaging Data Generation & Education (BRIDGE) Center using a Siemens MAGNETOM Prisma 3.0T research-dedicated MRI scanner and a 64-channel head/neck coil. Functional scans (T2-weighted) were run using a custom EPI multi-band pulse sequence (voxel size = 2 mm^3^ isotropic; number of slices = 69;gap size = 0 mm;matrix size = 106 x 106, FOV = 212 x 212 mm; TR = 2000 ms; TE = 30 ms;multiband acceleration factor = 3;phase encoding direction = A *>>* P, orientation = transversal;flip angle = 79^*°*^). To correct distortions in EPI sequences we used the TOPUP tool in the FMRIB Software Library (FSL; Jenkinson et al., 2012). We interleaved 2-3 pairs of TOPUP scans with functional runs within each session, with each pair consisting of forward and reverse phase-encoding directions. These scans were evenly distributed throughout the session. We also collected one high-resolution anatomical scan for each participant using a multi-echo MPRAGE sequence (voxel size = 1mm^3^ isotropic;number of slices = 176;matrix size = 256 x 256; TR = 2530 ms; TE = [1.69,3.55,5.41,7.27] ms; TI = 1240 ms;phase encoding direction = A *>>* P, orientation = sagittal;flip angle = 7°, GRAPPA acceleration factor = 2). This scan was used for segmentation, flattening, and delineation of functional ROIs on a reconstructed cortical surface.

### Task design for fMRI experiments

During scanning, participants viewed images targeting different visual regions. The experiment included two run types –NSD and BrainDiVE –which were interleaved within each session. Runs were identical in structure but differed in stimulus source: NSD runs used the top-activating natural images for each ROI, whereas BrainDiVE runs used images generated to maximally activate each ROI (see *Region-targeted images* for details).

Images were presented in a block design. Each block consisted of 12 images presented for 400 ms each with a 100 ms blank inter-stimulus interval, yielding a block duration of 6 s. Each block contained images from a single condition, corresponding to ROIs targeted by the stimuli: face-, body-, word-, place-, food-, FFA-, OFA-, OPA1-, and OPA2-targeted. A baseline condition (no images) was also included, for a total of 10 block types. Each block type was repeated six times per run in randomized order. Including additional blank periods at the start and end, each run lasted 384 s. Participants fixated on a central cross throughout the runs and performed a one-back task, pressing a button with their left index finger when two identical images appeared consecutively.

Each participant completed three NSD and three BrainDiVE runs in alternating order. The starting run type (NSD or BrainDiVE) was counterbalanced across participants. In the final set of participants (n = 12 after four participants were excluded), seven participants began with an NSD run.

In addition to the experimental runs, participants completed functional localizer runs to define category-selective visual regions (Jain et al., 2023; Stigliani et al., 2015). These runs were interleaved with NSD and BrainDiVE runs throughout the session. The localizer followed Jain et al., 2023, and included grayscale images from five categories (adult faces, bodies, houses, written words, and food), presented as isolated objects on phase-scrambled noise backgrounds (Jain et al., 2023). Images were presented in a block design (12 images per block, 400 ms per image, 100 ms gap), with each category repeated six times per run. Total run duration was 240 s. Participants fixated on a central cross and performed a one-back task (mean correct detection = 0.91 *±* 0.08;mean response time = 610.23 *±* 88.75 ms).

Stimuli were presented on a BOLDscreen32 LCD Display (Cambridge Research Systems, UK) and viewed via a mirror mounted on the head coil. Images were presented as squares (8°per side) on a mid-gray background.

### Region-targeted images

Region-targeted images were generated using BrainDiVE, which combines an image-computable voxel-wise encoding model with a latent diffusion model (LDM;Figure 1A). An overview of the method is provided here, complete details are described in Luo et al., 2023. We begin by constructing voxelwise encoding models, where the encoding model for the *i*-th voxel maps an image ℐ∈ℝ^*H×W ×*3^ to the corresponding predicted beta value 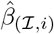. Our encoding models utilize pretrained deep neural networks (CLIP; Radford et al., 2021) as a backbone, followed by a learned per-voxel linear mapping (Naselaris et al., 2011; Wang et al., 2023). Once constructed, our models are end-to-end differentiable, which allows us to compute the gradient of a voxel’s predicted beta value with respect to the input image. The predicted beta value can be treated as an energy function *E*(ℐ), which allows us to view the gradient with respect to the input as the Stein score (i.e., score function) *s*(*x*) = Δ_ℐ_ log (*p*(ℐ)). However, directly optimizing images using the gradient yields degenerate images that are overfit to the encoding model. To prevent this, we leverage an image diffusion model to regularize the synthesis process (stable-diffusion-v-2-1-base; Rombach et al., 2022). During the reverse diffusion process, we guide the denoising using the encoding model prediction for a target visual region. At each step of denoising, we compute the predicted response of each voxel in the target ROI to the current estimated denoised image, and average over predicted voxelwise responses in the ROI. We then compute the gradient of this averaged response with respect to the diffusion latent, and perturb the diffusion latent in the direction of this gradient. We repeat this procedure for each denoising step. By additively combining the original diffusion model prediction with the encoder gradient, we are able to balance the naturalistic image objective and the predicted response maximization objective, resulting in naturalistic images that are predicted to activate a target brain region.

Using the above method, we generated images targeted to 9 different cortical regions identified in NSD participants (Allen et al., 2022). The first 5 regions were broad networks of category-selective voxels, identified by combining voxels from ROIs with shared category selectivity (face-selective regions: FFA, OFA, mTL-faces, aTL-faces;body-selective regions: EBA, FBA, mTL-bodies;word-selective regions: OWFA, VWFA, mfs-words, mTL-words;place-selective regions: OPA, PPA, RSC;food-selective regions: medial food-selective ROI, lateral food-selective ROI). Face, body, word, and place-selective regions were identified using an independent functional localizer (see Allen et al., 2022). Food-selective regions were identified using a mask of food-selective voxels created in previous work (Jain et al., 2023). This food-selective mask was created for each NSD participant individually, by using a general linear model to identify voxels significantly more responsive to food-containing NSD images versus other category labels, then intersecting this mask with an anatomical definition of the ventral visual cortex (Jain et al., 2023). We also generated images targeted to FFA and OFA individually, and for two disjoint clusters of voxels within the occipital place area (OPA), which were identified using von Mises Fisher (vMF) clustering on the encoding model linear mapping weights (Luo et al., 2023).

For each of these regions, we generated 1,000 images for each of the four NSD participants that completed all scanning sessions (S1, S2, S5, S7), giving a total of 4,000 images total for each target region, which were then ranked (see below) to give the final set of experimental stimuli. The BrainDiVE images were generated utilizing OpenCLIP laion2b s34b b88k ViT-B/16 as the frozen image-computable encoder backbone. We then trained a second voxelwise encoder for each of the four participants, using EVA-02 CLIP ViT-B/16 as a backbone. This second encoder was used to rank the images. The ranking encoder utilizes a different backbone to avoid potential adversarial examples –where an image is excessively overfit to activate a specific model. For all encoders, we trained them on the 9,000 unique images for each participant, using the final [CLS] embedding normalized to unit-norm. To rank images for each region, we combined all 4,000 images and ranked them for each participant using the second encoder’s predicted voxel activations (averaged across the voxels of interest). The ranks for each image were averaged across the four participants. For each region, the top-100 images with the lowest ranks were selected.

In addition to BrainDiVE images, we selected sets of natural images that were targeted to the same sets of voxels. Natural images were chosen from the actual stimulus images shown to participants in NSD (originally from the Microsoft COCO dataset; Lin et al., 2014). To select these images, we used the set of 1,000 images that were viewed by all NSD participants, and ranked these images based on the average response of the target ROI, for each of S1, S2, S5, and S7 separately. We then averaged the ranks across participants, and selected the top-100 images with lowest ranks (i.e., largest associated fMRI response).

### Preprocessing of fMRI data

MRI data preprocessing was performed using FSL (Jenkinson et al., 2012) and Freesurfer (available at: https://surfer.nmr.mgh.harvard.edu/). For each participant, we performed cortical surface gray-white matter volumetric segmentation of the high-resolution anatomical T1 scan using Freesurfer (recon-all; Dale et al., 1999). This segmented data was used to define functional ROIs and for surface-based analyses (see below). Functional preprocessing included unwarping (TOPUP distortion correction;see *Acquisition of fMRI data*), registration, motion correction, and detrending. Functional data from the first volume were aligned to the anatomical image using boundary-based registration for each participant (bbregister; Greve and Fischl, 2009), with the resulting transformation matrix applied to all functional runs. Motion correction was performed using FSL MCFLIRT (6 degrees of freedom, no spatial smoothing, and including a final sinc interpolation step; Jenkinson et al., 2002). Finally, temporal detrending was applied using a high-pass filter (1/100 Hz cutoff) to remove slow drifts.

### General linear modeling

Functional data from NSD, BrainDiVE, and functional localizer runs were analyzed using a general linear model (GLM; the FEAT FMRI Expert Analysis Tool in FSL). Each block type was treated as a separate condition and, for each, we convolved the stimulus onset with a canonical gamma hemodynamic response (phase = 0 s, SD = 3 s, lag = 6 s). These regressors were used to estimate both the response of each voxel to each condition (*β* values) and the contrast between conditions (*t*-statistics). GLMs were computed separately for individual runs. The GLM also included brain extraction (FSL Brain Extraction Tool; Smith, 2002) and pre-whitening to remove temporal autocorrelations (FILM; Woolrich et al., 2001). No additional motion correction, temporal filtering, or spatial smoothing was performed at this stage. Finally, for each participant, using a standard weighted fixed effects analysis, we separately combined data from the three NSD, BrainDiVE, and functional localizer runs. ROIs were defined using data from the functional localizer runs (see *Defining functional regions of interest*), and the *β* and *t* values from NSD and BrainDiVE runs were used in subsequent analyses.

### Defining functional regions of interest

Category-selective ROIs were identified in each participant by projecting *t*-statistics from the functional localizer onto an inflated cortical surface map and manually defining each region. Face-selective (fusiform face area/FFA, occipital face area/OFA), body-selective (extrastriate body area/EBA), word-selective (visual word form area/VWFA), and place-selective (parahippocampal place area/PPA, occipital place area/OPA) regions were identified using contrasts of each category versus the other four categories in the localizer task, following Stigliani et al., 2015. Food-selective regions (Bannert & Bartels, 2022; Jain et al., 2023; Khosla et al., 2022; Pennock et al., 2023) were identified using a contrast of food versus the other four categories following Jain et al., 2023. In most participants, two patches of food-selective voxels were identified on the ventral surface, located medial and lateral to the fusiform gyrus. These two patches were combined into a single ROI for our primary analyses (see Supplementary Table 1 for voxel counts). Following manual definition, each ROI was thresholded based on significance of the relevant contrast (*p <* 0.05;FDR corrected). For a small number of ROIs (OPA in S01, S07 and S09, and food-selective ROI in S08), this threshold yielded fewer than five voxels;in these cases, a more lenient threshold was applied (*t >* 2, uncorrected). This ensured that ROIs were defined in all participants, while retaining the most category-selective voxels where possible. Voxels overlapping across ROIs were excluded from our analyses. Final voxel counts are reported in Supplementary Table 1. For our primary analyses, left and right hemisphere ROIs were concatenated.

### Statistical analyses

All statistical analyses were performed in Python (version 3.13.3) using custom code, with functions from *SciPy* (version 1.16.0) and *statsmodels* (version 0.14.5). To test differences between experimental conditions and ROIs, we used repeated-measures ANOVAs, with *p*-values estimated via non-parametric permutation tests. For each of 10,000 permutations, values were shuffled within participants to preserve the repeated-measures structure, and *F*-statistics were computed for each effect. Final *p*-values were calculated as the proportion of permutations in which the shuffled *F*-statistic exceeded the observed *F*-statistic. This non-parametric method provides a more conservative alternative to standard parametric *p*-values for small sample sizes and follows analyses used in prior work (Henderson et al., 2022; Henderson et al., 2025; Rademaker et al., 2019; Sprague & Serences, 2013).

Paired comparisons between conditions (for ROI and behavioral data) were performed using permutation-based paired *t*-tests. For each of 10,000 permutations, condition labels were randomly swapped within participants (50% probability), and a *t*-statistic was computed for the shuffled data. Two-tailed *p*-values were calculated as twice the smaller of the proportions of permuted *t*-statistics greater than or less than the observed *t*-statistic.

### Surface-based distance estimation

To analyze how response properties varied across anatomical locations within the occipital place area (OPA), we estimated vertex positions along the anatomical posterior-anterior and medial-lateral axes. Voxel data were projected onto the cortical surface, and analyses were performed on surface mesh vertices. To ensure sufficient coverage of OPA, a more lenient threshold was used for defining the ROI in this analysis (*t >* 2, uncorrected). For each participant, posterior and medial anatomical landmarks were defined using parcels from the Destrieux atlas (Destrieux et al., 2010), obtained using Freesurfer’s recon-all. The occipital pole served as the posterior landmark and the parieto-occipital sulcus served as the medial landmark. These parcels were selected based on their locations posterior and medial to OPA;similar results were obtained using different but nearby parcels as landmarks. For each parcel, the median coordinate in volumetric space was used to define a single landmark vertex. For each surface vertex, geodesic distance to each landmark was computed using PyCortex (version 1.2.11; Gao et al., 2015), separately for each participant and hemisphere. These distances defined each vertex’s position along along the posterior-to-anterior (distance from posterior landmark) and medial-to-lateral (distance from medial landmark) axes.

### Stimulus image statistics

To quantify perceptual differences between image types, we computed a set of image statistics for each image used in the fMRI experiment. This included 100 NSD and 100 BrainDiVE images per ROI (see *Region-targeted images*), as well as two baseline sets: 1,000 randomly selected NSD images (125 images from each of 8 participants) and 1,000 random synthetic images generated using Stable Diffusion 2.1 base (no brain conditioning;0.75 SAG, DPM++solver).

To compute luminance and color statistics, images were converted from RGB to CIELAB color space, where L*is the luminance channel (0 to 100), a*is the green-red opponency channel (−127 to 127), and b*is the yellow-blue opponency channel (−127 to 127). For each image, we computed the mean of each channel and the standard deviation of L*as a measure of root-mean-square (RMS) contrast. We estimated color saturation by converting from RGB to HSV space and retaining the S channel.

Spectral properties were quantified using a Gabor filter bank (Henderson et al., 2023) with 8 spatial frequencies (logarithmically spaced from 4 to 72 cycles/stimulus;images were 256×256) and 12 orientations (0°-180°). For each frequency and orientation, a quadrature pair of filters (90°phase offset) was applied, and the image was convolved with each filter to obtain two feature maps (*a* and *b*). Response magnitude was then computed as 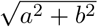 and averaged across the entire image.

Edge curvature was estimated using a bank of bent Gabor filters (Li & Bonner, 2020), with 24 orientations (0°-360°) and 6 bend values, all at 40 cycles/stimulus. Bend values were logarithmically spaced from 0 (linear) to 0.50 (~90°curve angle). Feature maps were computed using the same quadrature pair approach as for the standard Gabor filters, which resulted in a stack of feature maps corresponding to different bend and orientation values. Edge pixels were identified using the Roberts’Cross operator. For each edge pixel, the bend value corresponding to the maximal filter response was selected, and mean bend across edge pixels was used as the curvature estimate.

### Stimulus image behavioral evaluation

Human behavioral evaluation of model-generated images was conducted using an online experiment previously reported in Luo et al., 2023. Participants were recruited via Prolific (London, UK), and the experiment was implemented using the Gorilla Experiment Builder (Anwyl-Irvine et al., 2020). We recruited 10 participants for the OFA-FFA comparison and 10 participants for the OPA subregions comparison. The sample size was based on expected large effect sizes, with all analyses performed within-participant.

On each trial, participants viewed two groups of 10 images (arranged in a 2 x 5 grid) presented on the left and right sides of the screen. A prompt below the images showed a question (e.g., “Which group has more text?”), and participants responded with “Group 1”, “Group 2”, or “Same”. The display remained on the screen until the participant responded. Image comparisons were either OFA-versus FFA-targeted or OPA1-versus OPA2-targeted, using either NSD or BrainDiVE images. NSD and BrainDiVE images were not directly compared within trials.

The questions used in the OFA-FFA comparison were: “Which group…”

- “appears to have more human faces per image?”
- “has faces that are more photorealistic?”
- “has more abstract shapes and lines?”
- “has more animals or items visually similar to animals?”
- “has more food or items visually similar to food?”
- “has more text or items visually similar to text?”
- “has more weird faces (for example animal faces, low-quality faces, abstract faces, odd angles etc.)?”
- “looks more far away (as opposed to close-up)?”

The questions used in the OPA1–OPA2 comparison were: “Which group…”

- “looks more indoor (as opposed to outdoor)?”
- “looks more natural (as opposed to man-made)?”
- “looks more far away (as opposed to close-up)?”
- “has more angular/geometric lines?”
- “has more individual objects?”
- “has more people/animals?”

## Data and code availability

All source data and research code will be made publicly available upon acceptance of this manuscript.

## Acknowledgments

This work was partially supported by an award from Apple Inc to MJT. Any views, opinions, findings, and conclusions or recommendations expressed in this material are those of the author(s) and should not be interpreted as reflecting the views, policies or position, either expressed or implied, of Apple Inc.

## Supplementary Material

**Supplementary Figure 1:**
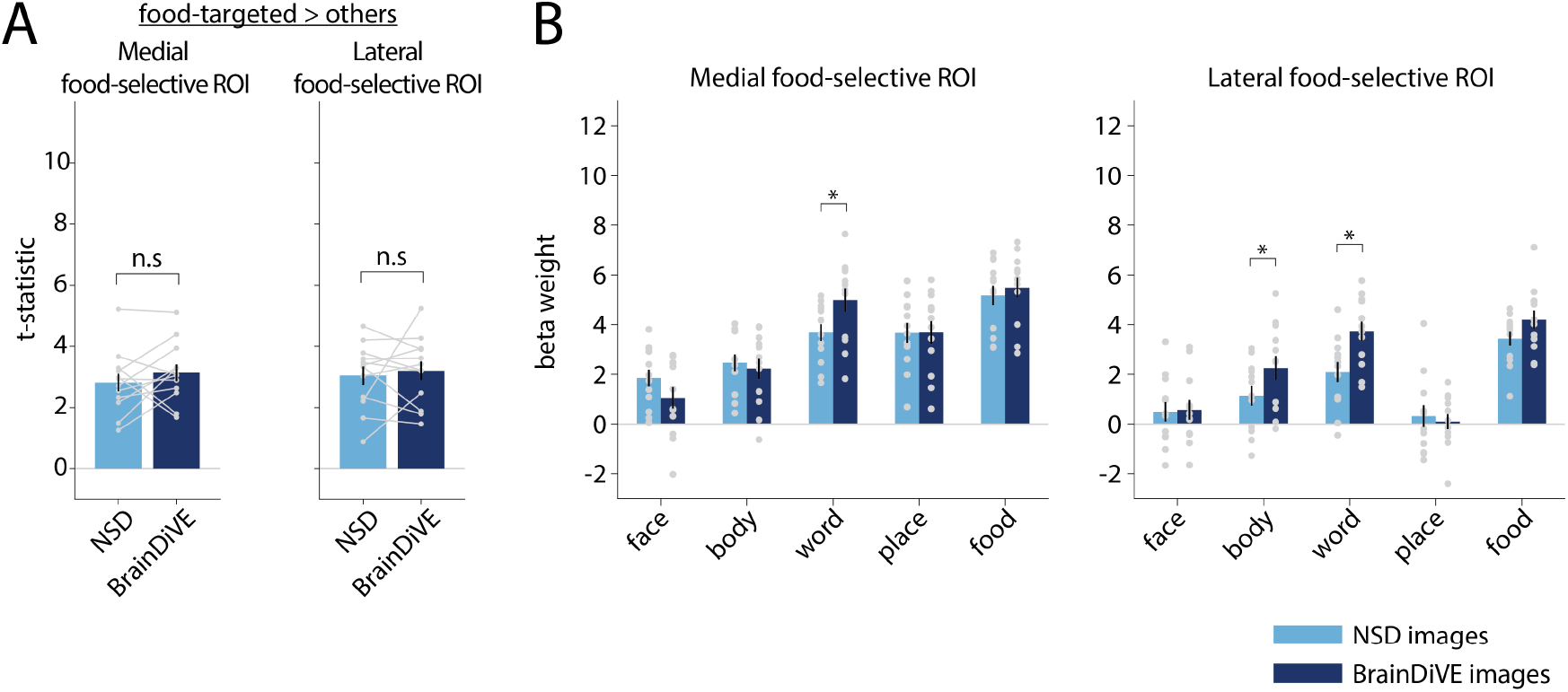
Responses elicited by BrainDiVE images and NSD images, for different subregions of food-selective ROI. **(A)** Average *t*-statistics for a contrast between images targeted for food-selective ROIs versus images that were targeted for other ROIs. Bar heights and error bars reflect the mean and SEM across participants (n=12), light gray dots and lines indicate single participant data, which is averaged across voxels within each ROI. **(B)** Beta weights associated with different image types (e.g., “face” refers to images targeted for face-selective ROIs in NSD participants). Different colors indicate beta weights from NSD or BrainDiVE runs. Bar heights and error bars indicate mean and SEM across participants (n=12);gray dots represent individual participants. In (A-B), brackets over pairs of bars indicate significance of the difference between NSD and BrainDiVE images (paired t-test with permutation, asterisk indicates p *<* 0.01, n.s. indicates p *>* 0.01).

**Supplementary Figure 2:**
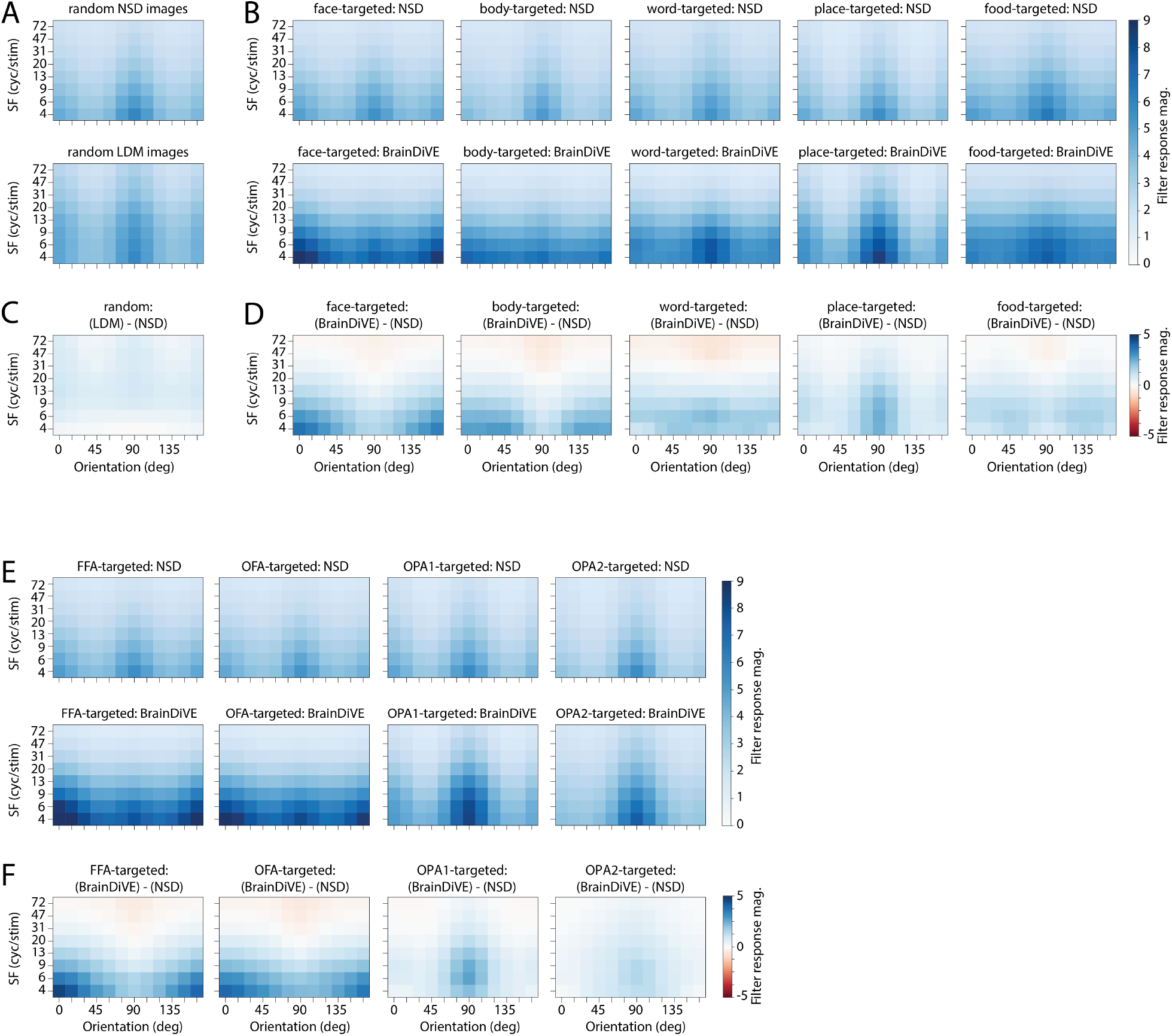
Differences in spectral statistics across image groups. For each image, average orientation and spatial frequency magnitude were estimated using a Gabor filter bank with 12 orientations and 8 spatial frequency values (see *Methods*). Each panel shows Gabor filter response magnitude, averaged across images, as a 2-dimensional heatmap. **(A)** Average filter response magnitude for 1000 randomly-selected NSD images (top) and 1000 LDM-generated images (bottom), synthesized using Stable Diffusion 2.1 base with no brain conditioning (see *Methods* for details). **(B)** Average filter response magnitude for region-targeted NSD and BrainDiVE images. **(C-D)** Difference between diffusion-generated and natural images in (A) and (B), respectively (i.e., bottom row of A-B minus top row). **(E-F)** similar to (B) and (D), shown separately for FFA-targeted, OFA-targeted, OPA1-targeted, and OPA2-targeted images.

**Supplementary Figure 3:**
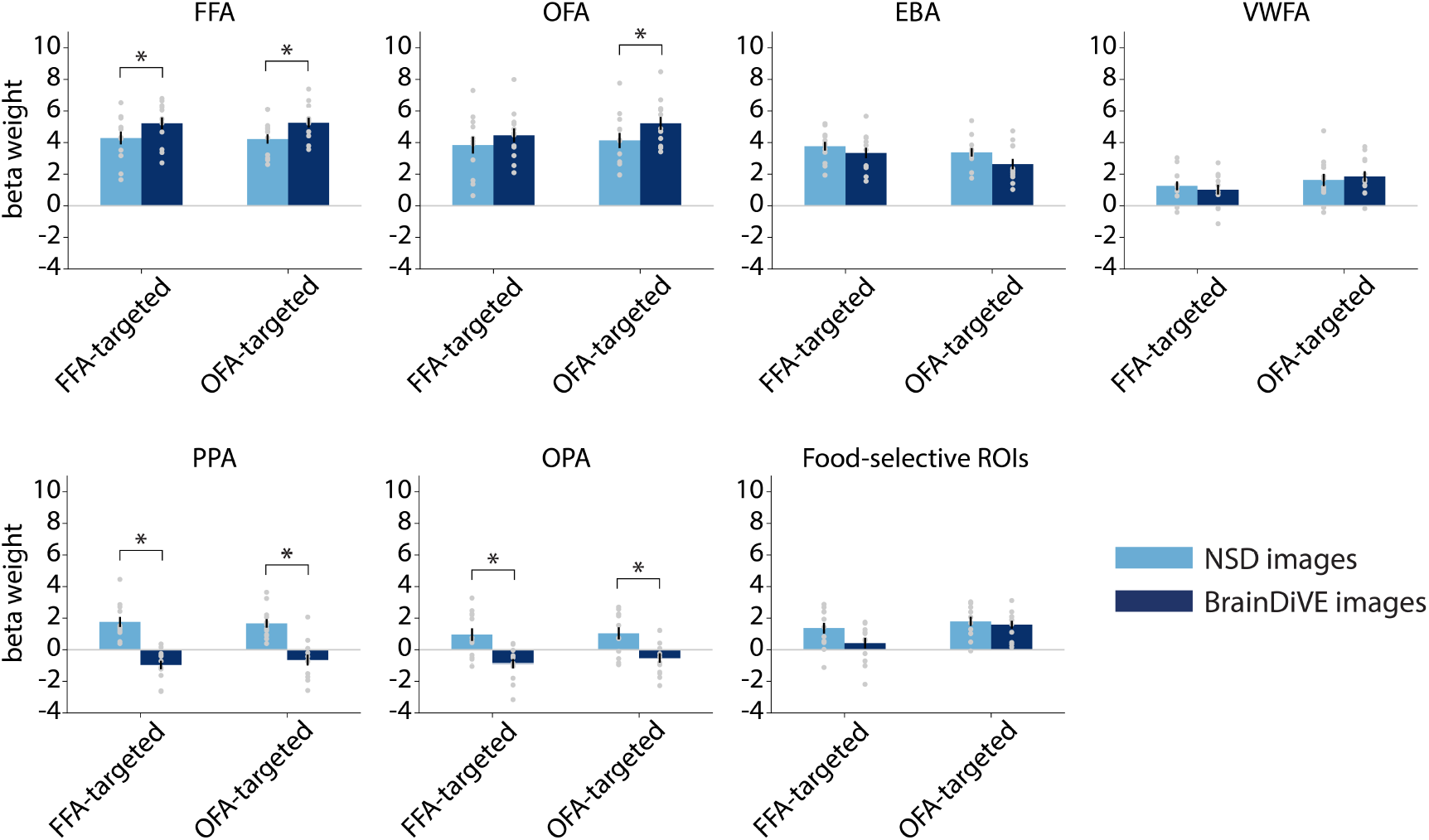
Responses elicited by OFA-targeted and FFA-targeted BrainDiVE images and NSD images, for the full set of ROIs. Each panel shows the *β* weights associated with each image type. Bar heights and error bars reflect the mean and SEM across participants (n=12), light gray dots indicate single participant data, which is averaged across voxels within each ROI. Brackets over pairs of bars indicate significance of the difference between NSD and BrainDiVE images (paired t-test with permutation;see *Methods*;asterisk indicates p *<* 0.01).

**Supplementary Figure 4:**
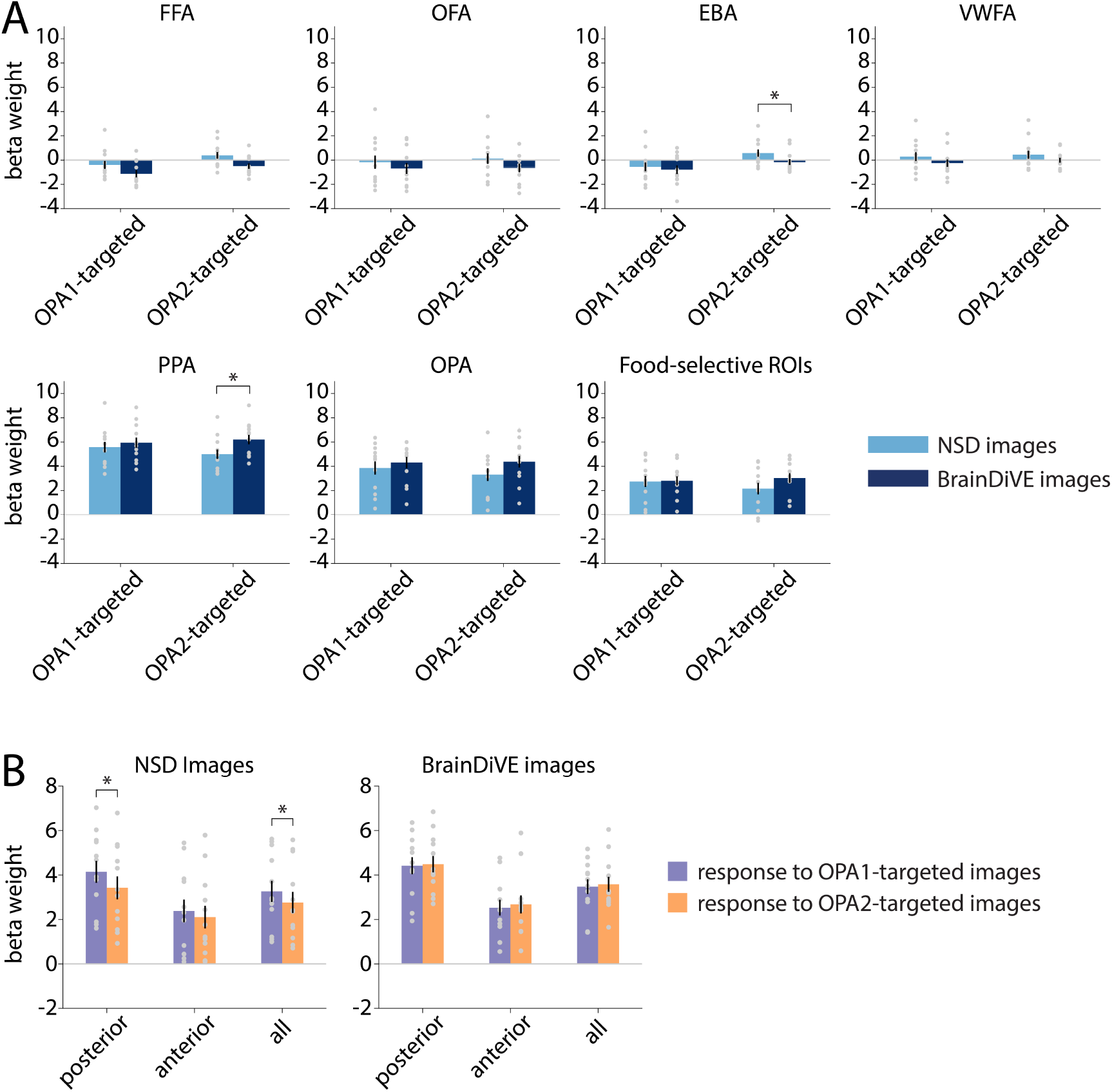
Responses elicited by BrainDiVE and NSD images targeted to different subregions of OPA (OPA1 = posterior subregion, OPA2 = anterior subregion). **(A)** Responses to OPA1-targeted and OPA2-targeted images from NSD versus BrainDiVE, for the full set of ROIs. **(B)** Response to OPA1-targeted and OPA2-targeted images, compared for posterior OPA vertices, anterior OPA vertices, and all vertices combined (all). Posterior and anterior OPA portions were defined using a median split on anatomical distance values;see *Methods* for details. This panel shows the same data as in Figure 7F, but with bars grouped differently. In (A) and (B), each panel shows the *β* weights associated with each image type. Bar heights and error bars reflect the mean and SEM across participants (n=12), light gray dots indicate single participant data, which is averaged across voxels within each ROI. Brackets over pairs of bars indicate significance of the difference between the pair of bars (paired t-test with permutation;see *Methods*;asterisk indicates p *<* 0.01).

**Supplementary Figure 5:**
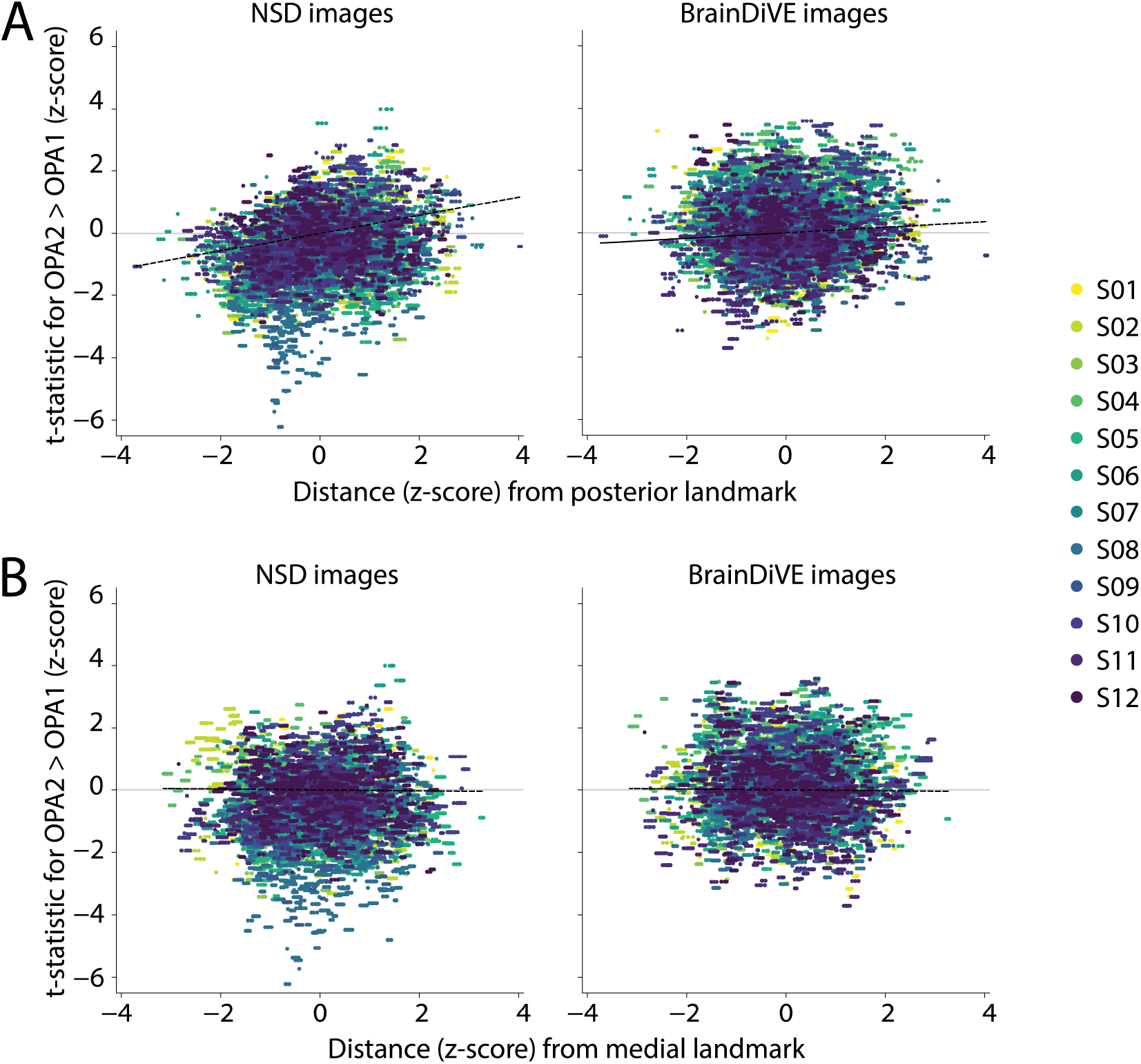
Anatomical organization of responses to OPA1-targeted and OPA2-targeted images. In each panel, the difference between response to OPA2-targeted vs. OPA1-targeted images (*t*-statistic, z-scored), is plotted as a function of anatomical position along a posterior-to-anterior **(A)** or medial-to-lateral **(B)** axis. Each dot is a vertex, colors indicate vertices from different participants. Dotted line indicates the best fit linear regression line.

**Supplementary Table 1.** Number of voxels in each functionally-defined ROI, following thresholding. See *Methods* for details on thresholding procedures. Voxel counts are shown for each hemisphere separately. We concatenated both hemispheres of each ROI for our main analyses.

|  | **S01** | **S02** | **S03** | **S04** | **S05** | **S06** | **S07** | **S08** | **S09** | **S10** | **S11** | **S12** |
| --- | --- | --- | --- | --- | --- | --- | --- | --- | --- | --- | --- | --- |
| **FFA, LH** | 15 | 25 | 145 | 255 | 31 | 325 | 27 | 105 | 90 | 43 | 199 | 107 |
| **FFA, RH** | 98 | 144 | 135 | 172 | 129 | 268 | 192 | 129 | 150 | 203 | 216 | 222 |
| **OFA, LH** | 24 | 5 | 28 | 126 | 14 | 24 | 9 | 30 | 9 | 0 | 118 | 52 |
| **OFA, RH** | 8 | 3 | 47 | 72 | 130 | 13 | 110 | 84 | 8 | 18 | 111 | 80 |
| **EBA, LH** | 538 | 136 | 326 | 73 | 285 | 372 | 75 | 84 | 138 | 359 | 154 | 324 |
| **EBA, RH** | 916 | 451 | 513 | 94 | 714 | 464 | 258 | 92 | 347 | 441 | 291 | 494 |
| **PPA, LH** | 14 | 24 | 54 | 142 | 20 | 109 | 12 | 67 | 5 | 48 | 127 | 106 |
| **PPA, RH** | 11 | 62 | 42 | 128 | 53 | 269 | 7 | 68 | 14 | 192 | 154 | 132 |
| **OPA, LH** | 148 | 19 | 20 | 45 | 38 | 80 | 45 | 32 | 95 | 70 | 40 | 113 |
| **OPA, RH** | 101 | 24 | 17 | 49 | 77 | 403 | 118 | 62 | 130 | 61 | 125 | 97 |
| **VWFA, LH** | 285 | 30 | 188 | 130 | 876 | 494 | 677 | 113 | 315 | 195 | 114 | 223 |
| **VWFA, RH** | 123 | 2 | 109 | 87 | 156 | 230 | 182 | 5 | 14 | 0 | 98 | 70 |
| **Food, LH** | 136 | 27 | 286 | 12 | 436 | 231 | 257 | 58 | 81 | 74 | 118 | 141 |
| **Food, RH** | 29 | 7 | 174 | 161 | 395 | 57 | 65 | 19 | 74 | 10 | 57 | 163 |

**Supplementary Table 2.** Test statistics comparing the response (beta weight) of each area to NSD images versus BrainDiVE images, targeted for different regions. See Figure 3 for the beta weight values on which these tests are performed. Each row shows the results of a two-sided paired *t*-test, with *p*-values obtained using a permutation test (see *Methods* for details). Positive *t* values indicate a larger response to BrainDiVE images versus NSD images of the specified type. Bolded *p*-values indicate values less than 0.01.

| ROI | Image Type | t | df | p |
| --- | --- | --- | --- | --- |
| FFA | face-targeted images | 5.88 | 11 | **0.0012** |
| FFA | body-targeted images | 1.10 | 11 | 0.2946 |
| FFA | word-targeted images | -2.11 | 11 | 0.0602 |
| FFA | place-targeted images | -1.59 | 11 | 0.1462 |
| FFA | food-targeted images | -2.88 | 11 | 0.0146 |
| OFA | face-targeted images | 3.32 | 11 | **0.0096** |
| OFA | body-targeted images | 0.69 | 11 | 0.5118 |
| OFA | word-targeted images | 0.57 | 11 | 0.5734 |
| OFA | place-targeted images | -1.79 | 11 | 0.1074 |
| OFA | food-targeted images | -1.10 | 11 | 0.2944 |
| EBA | face-targeted images | 0.42 | 11 | 0.6814 |
| EBA | body-targeted images | 3.60 | 11 | **0.0050** |
| EBA | word-targeted images | -3.41 | 11 | **0.0076** |
| EBA | place-targeted images | -0.50 | 11 | 0.6248 |
| EBA | food-targeted images | -2.06 | 11 | 0.0640 |
| VWFA | face-targeted images | 0.74 | 11 | 0.4846 |
| VWFA | body-targeted images | 1.19 | 11 | 0.2582 |
| VWFA | word-targeted images | 6.04 | 11 | **0.0008** |
| VWFA | place-targeted images | -1.68 | 11 | 0.1128 |
| VWFA | food-targeted images | -0.13 | 11 | 0.8862 |
| PPA | face-targeted images | -8.09 | 11 | **0.0006** |
| PPA | body-targeted images | -3.76 | 11 | **0.0024** |
| PPA | word-targeted images | -3.15 | 11 | **0.0088** |
| PPA | place-targeted images | 2.51 | 11 | 0.0306 |
| PPA | food-targeted images | -4.40 | 11 | **0.0022** |
| OPA | face-targeted images | -3.27 | 11 | **0.0086** |
| OPA | body-targeted images | -1.85 | 11 | 0.0762 |
| OPA | word-targeted images | -0.32 | 11 | 0.7738 |
| OPA | place-targeted images | 1.60 | 11 | 0.1380 |
| OPA | food-targeted images | -2.01 | 11 | 0.0750 |
| Food | face-targeted images | -1.17 | 11 | 0.2846 |
| Food | body-targeted images | 1.55 | 11 | 0.1518 |
| Food | word-targeted images | 5.55 | 11 | **0.0002** |
| Food | place-targeted images | -0.17 | 11 | 0.8564 |
| Food | food-targeted images | 1.76 | 11 | 0.0502 |

